# A method to estimate the effective point spread function of static single molecule localization microscopy images

**DOI:** 10.1101/2022.03.05.483117

**Authors:** Thomas R. Shaw, Frank J. Fazekas, Sumin Kim, Jennifer C. Flanagan-Natoli, Emily R. Sumrall, S. L. Veatch

**Affiliations:** Program in Biophysics, University of Michigan, Ann Arbor, Michigan; Program in Applied Physics, University of Michigan, Ann Arbor, Michigan; Program in Cellular and Molecular Biology, University of Michigan, Ann Arbor, Michigan

## Abstract

Single molecule localization microscopy (SMLM) permits the visualization of cellular structures an order of magnitude smaller than the diffraction limit of visible light, and an accurate, objective evaluation of the resolution of an SMLM dataset is an essential aspect of the image processing and analysis pipeline. Here we present a simple method that uses the pair autocorrelation function evaluated both in space and time to measure the time-interval dependent effective point spread function of SMLM images of static samples. Using this approach, we demonstrate that experimentally obtained images typically have effective point spread functions that are broader than expected from the localization precision alone, due to additional uncertainty arising from factors such as drift and drift correction algorithms. The method is demonstrated on simulated localizations, DNA origami rulers, and cellular structures labelled by dye-conjugated antibodies, DNA-PAINT, or fluorescent fusion proteins.

**STATEMENT OF SIGNIFICANCE:** Single molecule localization microscopy (SMLM) is a class of imaging methods that resolve fluorescently labeled structures beyond the optical resolution limit of visible light. SMLM detects stochastically blinking labels over time, and localizes each blink with precision of order 10 nm. The effective resolution depends on factors such as signal-to-noise ratio, localization algorithm, and several post-processing steps such as stage drift correction. We present a method to evaluate this effective resolution by taking advantage of temporal correlations of fluorophore blinking to separate the distribution of pairs of localizations from the same molecule from those from different molecules. The method is robust on useful timescales for a range of SMLM probes.

## INTRODUCTION

Localization microscopy is a powerful tool to image structures in cells with dimensions ranging between tens of nanometers to tens of microns. Methods such as (d)STORM (1, 2), (F)PALM (3, 4), and PAINT (5) exploit the stochastic blinking of single fluorophores to localize emitting molecules with a localization precision much smaller than the diffraction limit of visible light, by imaging only a small subset of probes in any given image frame. These samples are then imaged over time, and acquired localizations are typically assembled into a single reconstructed super-resolved image.

Assessing the quality of reconstructed images can be challenging as numerous factors can contribute. These factors can include the labeling density, the brightness and blinking dynamics of the fluorophore, motions of the stage or labeled molecules during acquisition, and the analytical methods used in post-processing. One important measure of the quality of a measurement is the localization precision of single fluorophores, which is influenced by many of the factors listed above. Many localization algorithms directly return the localization precision of single fits and similar information can be extracted directly from the localizations themselves through the use of pair-correlation functions or nearest neighbor analyses that extract the distribution of positions of molecules detected in adjacent frames (6, 7). Other metrics of image quality have been developed that integrate both precision and spatial sampling, reporting superior resolution when localizations effectively sample labeled objects (8, 9). One widely used method, called Fourier Ring Correlation (FRC) (10–13), effectively captures the impact of factors that degrade quality over the entire span of image acquisition, such as sample drift and imperfect drift correction, or the recently-reported observation of. A disadvantage of FRC on reconstructed super-resolution images is that the absolute number produced is not directly related to how accurately single molecule positions are be measured. The absolute value of the FRC depends on the types of structures imaged, how well they are sampled, and the specific regions of interest used.

Here we present a simple method to estimate the effective point spread function (PSF) of a localization dataset as a way to evaluate its resolution. The estimate incorporates errors relevant to measuring distances between labeled objects within images. This resolution metric is expected to be the most useful when the goal of the imaging experiment is to measure distances between labeled components, for example when constraining the structure of a multi-protein complex or when tabulating the local density of labeled components. The method presented exploits temporal correlations in the blinking dynamics of single fluorophores commonly used for localization microscopy (14–16). This enables the method to report on errors accumulated over time that do not typically impact the localization precision but impact how accurately a molecule’s position is determined relative to others. Here, we derive the proposed resolution metric, validate it through simulation and an image of DNA origami rulers (17), and apply it to several images of labeled structures in cells.

## MATERIALS AND METHODS

### Simulations

Simulations mimicking DNA origami rods were accomplished by randomly placing pairs of fluorophores positioned 50nm apart within a 40μm by 40μm region of interest at an average density of 1 pair per μm^2^. 20,000 individual image frames with an effective frame time of 0.1 sec were simulated by sampling a subset of molecular positions with a localization precision of lOnm in each lateral dimension. The dynamics of individual fluorophores were governed by a continuous time Markov process involving five states: one on state (1), three dark states (0, 01, 02), and an irreversible bleached state (B), following the procedure described previously (16, 18). The on state was accessible from any of the dark states, while dark state 0 was accessible only from the on state, and dark states 01 and 02 were accessible only from the previous dark states, 0 and 01, respectively. We used the following parameters (using the notation described in (16, 18)) to capture essential elements of our experimental observations: *λ*(0 → 1) = 1.2 Hz; *λ*(0 → 01) =0.05 Hz; *λ*(01 → 02) =0.0033 Hz; *λ*(01 → 1) =0.02 Hz; *λ*(02 → 1) =0.0005 Hz; 2(1 → 0) = 5 Hz; μ(1 → *B*) =0.05 Hz. The continuous time Markov process was simulated with the MATLAB File Exchange code “simCTMC.m” (19). When present, drift was applied to all molecular positions with a constant rate of 0.3nm/sec in the x direction along with diffusive drift characterized by a diffusion coefficient of D=2.5nm^2^/sec. Drift was corrected using the mean-shift algorithm described previously (20) using 1000 frames per alignment (10s).

### Experimental sample preparation

DNA origami “gatta-STORM” nanorulers were purchased from Gattaquant GMBH (Grafelfing, Germany) and a sample was prepared following manufacturer’s instructions. Briefly, biotinylated Bovine Serum Albumin (BSA; ThermoFisher; 1 mg/ml) was absorbed to a clean 35mm #1.5 glass-bottom dish (MatTek well; MatTek Life Sciences) for 5min then washed. Streptavidin was then applied (1 mg/ml) for 5min, then washed with a solution of phosphate buffered saline (PBS) plus 10mM MgCl_2_. A solution containing the biotinylated DNA origami was then applied. Samples were then washed and imaged in an imaging buffer supplemented with 10mM MgCl_2_. “gatta-PAINT” 80RG nanorulers in a sealed sample chamber were purchased from Gattaquant GMBH and imaged in the Atto655 color channel following manufacturer recommendations.

Mouse primary neurons were isolated from P0 mouse pups as described previously and cultured on MatTek wells (20). On day 10 of culture (days *in vitro* 10), neurons were rinsed with sterile Hank’s Balanced Salt Solution and fixed for 10min with pre-warmed 4% PFA (Electron Microscopy Sciences) in Phosphate Buffered Saline (PBS). The fixed neurons were rinsed three times with PBS and permeabilized in 0.2% Triton X-100 (Millipore Sigma) in PBS for 5min. Neurons were then incubated in blocking buffer containing 5% BSA for 30min, and labeled with Nup210 polyclonal antibody diluted in PBS (1:200; Bethyl laboratories A301-795A) overnight in 4□3°C. The following day, neurons were washed three times in PBS and stained with goat-anti-rabbit AlexaFluor 647 Fab Fragment (1:800; Jackson ImmunoResearch 111-607-003) for an hour, washed three times with PBS, then imaged.

CH27 B-cells (mouse, Millipore Cat# SCC115, RRID:CVCL_7178), a lymphoma-derived cell line (21) were acquired from Neetu Gupta (Cleveland Clinic). CH27 Cells were maintained in culture as previously described (22). Cells were adhered to MatTek wells coated with VCAM following procedures described previously (23). Briefly 0.1 mg/ml IgG, Fcγ-specific was adsorbed to a plasma cleaned well for 30 min at room temperature. Wells were rinsed with PBS, then nonspecific binding was blocked with 2% BSA at room temperature for 10 minutes, followed by incubation with 0.01 mg/mL recombinant human VCAM-1/CD106 Fc chimera protein (R&D Systems) and 0.01 mg/mL ChromPure Human IgG, Fc fragment (Jackson Immunoresearch) for 1 hour at room temperature or overnight at 4°C. VCAM-1 coated dishes were stored up to 1 week in VCAM-1 and Fc at 4°C. Immediately prior to plating, dishes were blocked at room temperature in 2% goat serum (Gibco) for 10 min, then cells were allowed to adhere for 15 min in media prior to chemical fixation in 2% PFA and 0.2% glutaraldehyde (Electron Microscopy Sciences). FActin was stained by permeabilizing cells with 0.1% Triton-X-100 prior to incubation with 3.3 μM phalloidin-AlexaFluor647 (Invitrogen) for at least 15 min. Phalloidin stained cells were imaged immediately after removing label. Cells transiently expressing Clathrin-GFP were permeabilized after fixation with 0.1% Triton-X-100 followed by labeling with a single domain anti GFP antibody (MASSIVE-SDAB 1-PLEX) from Massive Photomics GMBH (Grafelfing, Germany) for 1h at room temperature, then imaged in 0.5nM of imaging strand in the imaging buffer supplied by the manufacturer.

Cells expressing the membrane label Src15-mEos3.2 were prepared by transiently transfecting 10^6^ cells with a 1 μg of plasmid encoding Src15-mEos3.2 (N’-<u>MGSSKSKPKDPSQRR</u>NNNNGPVAT-rmEos3.21-C’) which was derived from a GFP tagged version by replacing GFP with mEos3.2 (24, 25). Transfection was accomplished by Lonza Nucleofector electroporation (Lonza, Basel, Switzerland) with program CA-137 and cells were grown in flasks overnight prior to plating and fixation as described above.

### Single molecule imaging and localization

Imaging was performed using an Olympus IX83-XDC inverted microscope. TIRF laser angles where achieved using a 100X UAPO TIRF objective (NA = 1.50), and active Z-drift correction (ZDC) (Olympus America). AlexaFluor647 was excited using a 647 nm solid state laser (OBIS, 150 mW, Coherent) and mEos3.2 was excited using a 561nm solid state laser (Sapphire 561 LP, Coherent), both coupled in free-space through the back aperture of the microscope. Fluorescence emission was detected on an EMCCD camera (Ultra 897, Andor). Samples containing AlexaFluor647 were imaged in a buffer containing 100mM Tris, 10mM NaCl, 550mM glucose, 1% (v/v) beta-mercaptoethanol, 500 μg/ml glucose oxidase (Sigma) and 40 μg/ml catalase (Sigma), with lOmM MgCl_2_ for the DNA origami sample. Samples with mEos3.2 or DNA PAINT Atto655 probes were imaged in imaging buffer from Massive Photomics GMBH. Single molecule positions were localized in individual image frames using custom software written in Matlab. Peaks were segmented using a standard wavelet algorithm (26) and segmented peaks were then fit as single emitters on GPUs using previously described algorithms for 2D (27), or as multi-emitters on a CPU using the ThunderStorm ImageJ plugin (28). After localization, points were culled to remove outliers prior to drift correction (20). Images were rendered by generating 2D histograms from localizations followed by convolution with a Gaussian for display purposes. For the nano-ruler samples, localizations were assigned to single fluorophores using a home-built implementation of DBSCAN (29), with ò = 12 nm and minPts = 15.

### Evaluation of space-time autocorrelations

Space-time autocorrelations were tabulated by first tabulating space- and time-displacements between all pairs of localizations within a specified region of interest (ROI) detected in a given dataset. This was accomplished using a *crosspairs()* function based on the one from the R package spatstat (1), but used here as a C routine with a MATLAB interface, as described previously (20). Lists of displacements were converted into space-time autocorrelation functions by binning in both time and space within the C routine for improved performance, followed by a normalization implemented in Matlab that produces a value *g*(*r, τ*) = 1 when localizations are randomly distributed in both space and time within the specified ROI. A derivation of the form of this normalization and an explanation of how it is computed is presented in Supplementary Note 1.

### Estimation of *g_PSF_* (*r, τ*) and *σ_xy_*(*τ*)

The core computations of the effective PSF estimation are gathered in a single MATLAB function. First, *g*(*r_i_, τ_j_*) is computed as described above, for a range of distance and time separation values. By default, *r_i_, i* = 1,…, *N_r_* represent bins with bin edges from 0 to 250 with equal spacing of 5nm, resulting in bin centers ranging between 2.5 to 247.5nm, and *τ_j_, j* = 1,…, *N_τ_* represent bins with edges that are log-spaced. The lower edge of the final time-separation bin is determined by identifying the lowest *τ* that satisfies 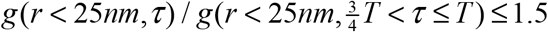, where T is the time of the image acquisition. The *τ*_max_ reported in figures is the bin center of this final time-separation bin.

Then, Δ*g*(*r,τ_j_*) = *g*(*r, τ_j_*) – *g*(*r,τ_max_*) are computed fo reach *τ*, normalized by their first spatial points (*r < 5nm*), and fitted to a Gaussian of the form 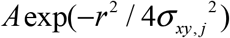, using MATLAB’s nonlinear least squares fitting routing fit(). *σ_xy,j_* is reported as the estimate of *σ_xy,j_* (*τ_j_*). Bootstrapped standard errors are determined by choosing 8 subsamples of the points, each containing ¼ as many points as the full dataset, and estimating *σ_xy_*(*τ_j_*)_*k*_ for each subsample *k*, in the same way as for the full dataset. The standard error is reported as 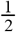 *std.dev*.(*σ_xy_*(*τ_j_*)_*k*_), where the 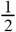 accounts for the overestimate of errors due to using 4 times fewer points.

### Measuring *g_PSF_*(*r,τ*) in simulations by grouping localizations with molecules

In simulations, localizations imaged at *x_i_, y_i_, τ_i_* are associated with the molecules that produced them. We tabulate displacements between all pairs associated with the same molecule 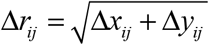 and Δ*t_ij_*. The list of all pairs is binned into two dimensional histograms following the same r and τ bin-edges as described for computing *g*(*r,τ*) above and are normalized by the number of pairs contributing to each bin. Distributions at each τ bin are fit to the same Gaussian form as applied to the *g_PSF_*(*r,τ*) estimated from Δ*g*(*r, τ*).

### Determining the resolution with Fourier Ring Correlation

The resolution of each dataset was assessed with Fourier Ring Correlation (FRC) (12). To produce the FRC curves, localizations were divided into consecutive blocks of 500 frames, and these blocks were randomly placed into one of two statistically independent subsets. For the simulated and experimental DNA origami datasets, as well as the nuclear pore complex dataset, the pixel size for the FRC calculation was taken to be 5nm, and square regions 10μm on a side were used as a mask. For the actin and src15 datasets, the pixel size was 10nm, with the mask 20μm on each side. 20 randomly determined repetitions of the calculation were performed for each dataset.

## RESULTS

### Derivation of the effective PSF estimate

The spatial autocorrelation function describing a distribution of static molecules is given by *g_molecules_*(*r*) and is tabulated as described in Methods and Supplementary Materials. This function can be divided into two components:

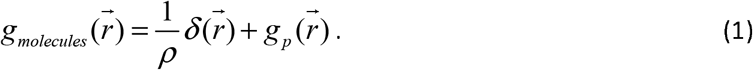

The first term in Eqn. 1 comes from counting single emitters and is a Dirac delta function 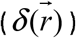 with magnitude equal to the inverse average density of molecules (*ρ*) over the region of interest (ROI). The second term in Eqn. 1 comes from correlations between distinct pairs of molecules and reports on the detailed structure present in the image. In SMLM, single emitters labeling molecules have dynamics governed by the probe photophysics, which can be described with the temporal autocorrelation function *g_e_*(*τ*). Probes can remain on for multiple sequential image frames and can blink on again at a later time before eventually bleaching irreversibly (14–16). As a result, *g_e_*(*τ*) is highly correlated (>1) at short time-intervals and decays sharply on time-scales describing the average on-time of fluorophores. This function continues to decay slowly at long τ, both because some probes tend to flicker over medium to long time-scales and because some fluorophores eventually bleach. Including *g_e_*(*τ*) produces the following spatio-temporal autocorrelation function for the emitting molecules:

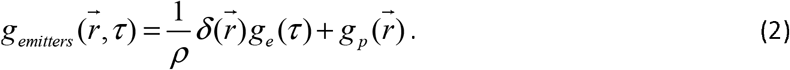

Eqn. 2 assumes the blinking statistics of different fluorophores are uncorrelated, which is why *g_e_*(*τ*) multiplies only the first term.

When fluorophores are observed with finite spatial resolution, the observed autocorrelation is blurred (convolved) by the autocorrelation function that describes that resolution, which in microscopy is typically called the effective PSF, or 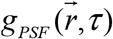. Including this factor, the autocorrelation function for the observed image, 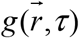, becomes:

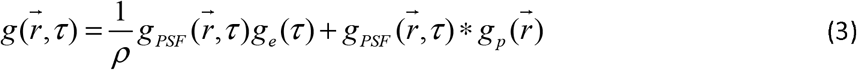

where * indicates a convolution. The first term in Eqn. 3 describes multiple observations of the same molecule and contains the most information on the PSF of the measurement. To isolate this component, we make several approximations and compare correlation functions tabulated at from pairs observed at time interval τ to those tabulated from pairs observed at large time separations *τ* = *τ*_max_ where the value of *g_e_*(*τ*)is relatively small.

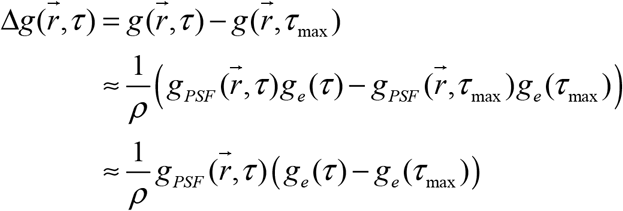

Where an appropriate choice of *τ_max_* □ *τ* depends on the duration of acquisition and on the blinking dynamics that are present in a given dataset. More simply written:

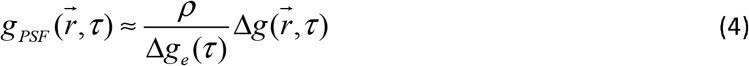

In words, Eqn. 4 states that the shape of the effective PSF can be estimated by comparing autocorrelation functions tabulated at different time-intervals. This observation forms the basis for the method demonstrated in this report.

To convert 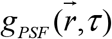 to a resolution length-scale, we assume that it takes on a Gaussian form:

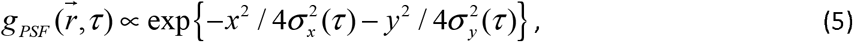

where *σ_x_*(*τ*) is the standard deviation in the *x* direction of the distance between the true position of the molecule at time *t* and the and a localization at time *t + τ*. The extra factor of 2 in the denominator accounts for the fact that *g_PSF_* reports on the distribution of distances between pairs of localizations, resulting in twice the variance compared to the error in one localization. Typically the effective PSF is isotropic in the lateral dimensions so we take *σ_xy_*:= *σ_x_* = *σ_y_* and compute angularly averaged correlation functions resulting in:

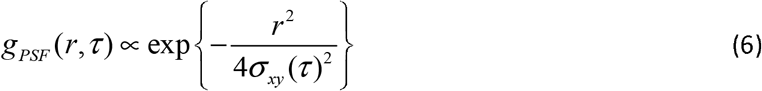

It is convenient to also define the mean squared displacement 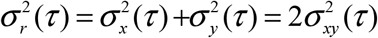 to evaluate localization precision, which accounts for errors in both dimensions. When localizations are acquired in three dimensions, the axial resolution often differs and this component can be considered independently:

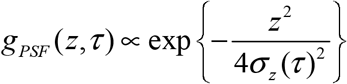

Two approximations are made in the process of arriving at Eqn. 4 that introduce practical limitations to this approach. The first and more minor limitation is that 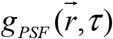 must not degrade too rapidly in *τ*. This assumption enables the cancelation of the term involving 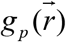 which captures image-specific factors that otherwise complicate the measurement of 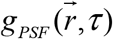. In practice, this assumption is largely valid since SMLM images are typically corrected for long time-scale errors through drift correction algorithms. Moreover, the amplitude of 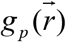 is typically much smaller than 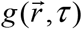 at short 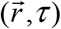 where this method is most useful, therefore errors in this subtraction have a minor impact on the result overall.

A second assumption required to arrive at Eqn. 4 provides a more direct practical limitation to the application of this method. This expression approximates 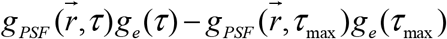 as 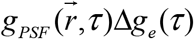, which is valid when *g_e_*(*τ*) ≫ *g_e_*(*τ*_max_) or when 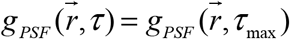. In essence, this approximation dictates that the method can be simply applied only up to time-scales where meaningful temporal correlations are observed for single fluorescent molecules, since correlated observations of localizations from the same particles are used to evaluate resolution. Past studies document long correlation times for many (d)STORM and (f)PALM probes (14–16) under a range of imaging conditions, supporting that this assumption will be valid over a broad range of time-intervals. PAINT probes, which produce localizations through binding and unbinding of a probe fluorophore to a target molecule, are not expected to produce temporal correlations that extend far beyond the off-rate of the specific binding interaction, since the binding of new probes from solution does not depend on the history of probe binding to a specific site (5, 30). For these probes, Eqn. 4 will only produce accurate results over time-scales relevant to probe un-binding. Plots of *g*(*r* < 25nm, *τ*) capture *g_e_*(*τ*) up to a numerical offset, and examples showing this decay for several experiments with different fluorophores are shown in Supplementary Figure S1.

Outside of the regime where *g_e_*(*τ*) ≫ *g_e_*(*τ*_max_), the subtraction of 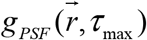 can lead to systematic errors in the determination of the width of 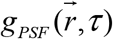 from 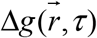, as demonstrated in Supplementary Figure S2. For cases where 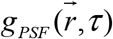 broadens slightly over time, for example when labeled molecules diffuse over length-scales comparable to the localization precision over the acquisition time, Eqn. 4 will produce a systematically narrow estimate of 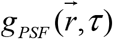 for *g_e_*(*τ*) □ *g_e_*(*τ_max_*), as demonstrated on simulated data in Supplementary Figure S3. A user could extend the applicability of this method to larger τ through a more complicated analysis to extract 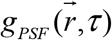 and 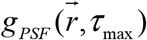 independently, or could tabulate 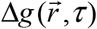 for closely spaced τ where changes in 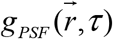 are expected to be more subtle. For the purposes of this report, we do not plot estimates of 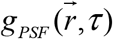 that may be subject to this systematic bias, using a cutoff of Δ*g*(*r* < 5nm, *τ*)/*g*(*r* < 5nm, *τ*_max_) < 0.5. In practice, we find that 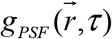 remains nearly constant overtime-scales larger than those used for drift-correction, so we apply the *σ_xy_* evaluated for the final accessible temporal bin to describe the localization precision of the remaining pairs.

The decay of *g_e_*(*τ*) means that the signal to noise ratio of 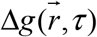 will degrade since fewer correlated pairs are observed at longer time-intervals. To increase statistical significance, we group time-intervals into increasingly large disjoint τ bins and estimate *g*(*r, τ*) as a weighted average over the tau bin containing τ. We typically report the central τ value of the bin, but occasionally show the full range of τ values included in the bin, for clarity. The τ bin edges are log-spaced to account for the exponential decay inherent in *g_e_*(*τ*),and with 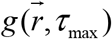 encompassing all τ greater than the cutoff indicated above, or the back quarter of the dataset 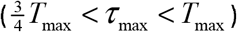, whichever is greater. Statistical confidence is estimated through bootstrapping and we estimate the statistical power of 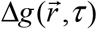 directly from 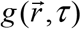, as described in Materials and Methods.

We have implemented this method as MATLAB code, which is available online(31).

### Validation through Simulation

To validate this approach, we generated simulated datasets of DNA origami nanorulers in which fluorophores are separated by a fixed distance of 50 nm. The blinking of the fluorophores were subject to a photophysical model based on (16,18). Briefly, fluorophores could exist in an “on” state, one of three dark states, or a bleached state. Transitions between states were governed by a continuous time Markov process, with transition rates roughly based on those measured in (16) but modified to reflect the experimental conditions used to obtain experimental images in this work. Nanorulers were placed randomly and uniformly with an average density of 1/μm^2^ across a 40μm by 40μm field of view with the molecules having a localization precision of 10 nm in each lateral dimension. 20,000 image frames were simulated with a frame time of 0.1 s. Figure 1a illustrates a small field of view containing 3 nanorulers, both as a reconstructed image and with localizations colored by time. An image showing a larger subset of the field of view is shown as Supplementary Figure S4.

**Figure 1:**
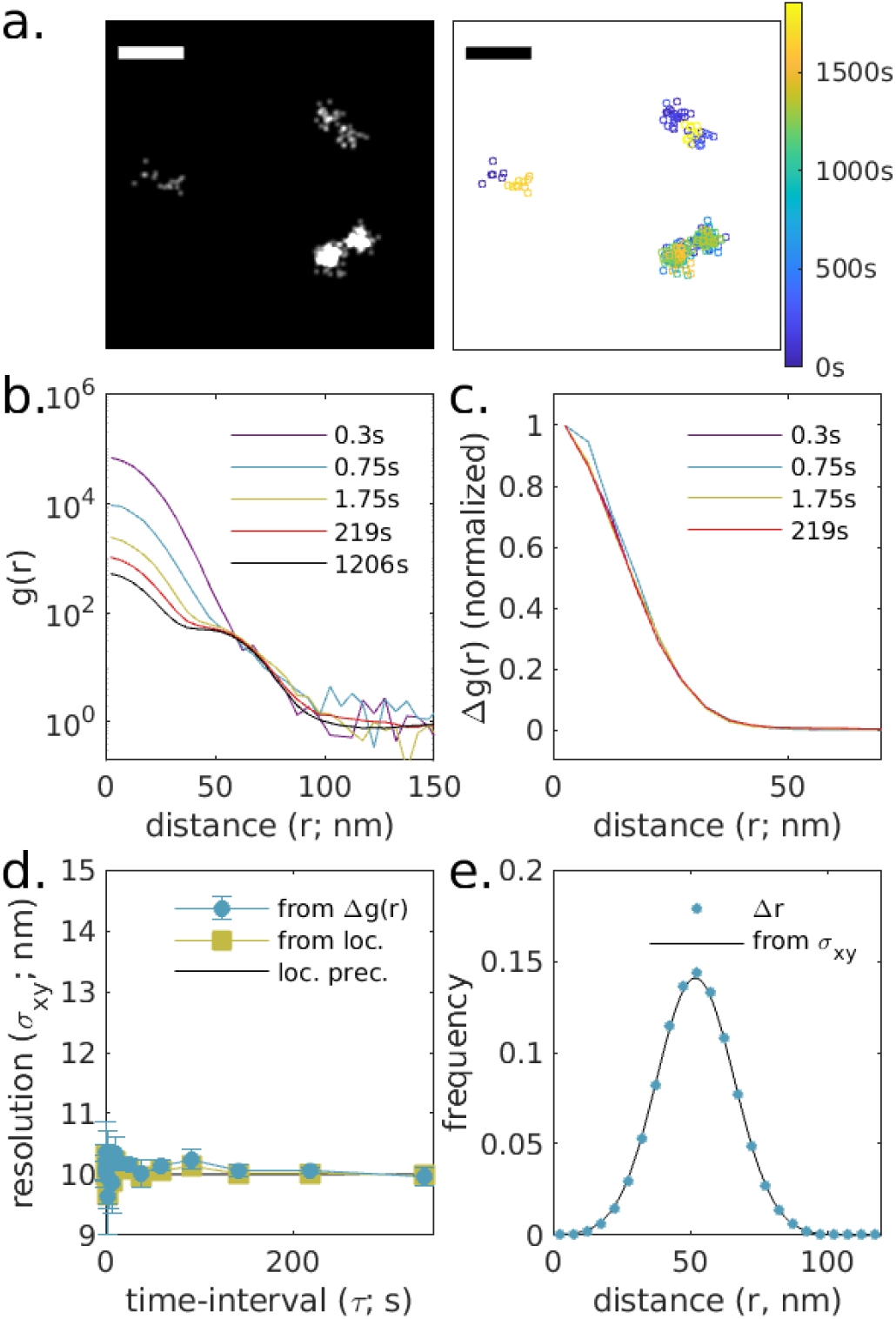
Validation of approach through simulation, (a.) Simulations consist of randomly positioned pairs of molecules positioned 5Onm apart in random orientations. Reconstructed image (left) and scatterplot of localizations with color representing the observation time (right) for a small subset of the simulated plane. Scale bar is 100nm. A reconstructed image showing a larger field of view is shown in Supplementary Figure S4. (b.) Auto-correlations as a function of displacement, *g*(*r, τ*), tabulated from simulations for time-interval windows centered at the values shown. (c.) Δ*g*(*r, τ*) = *g*(*r, τ*) – *g*(*r, τ*_max_ = 1206*s*)for the examples shown in b. (d.) Δ*g*(*r, τ*) are fit to 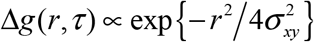 to extract the width of the effective point spread function in each lateral dimension, which in this case is the same as the resolution deduced by grouping localizations with their associated molecules (from loc.) and the localization precision (loc. prec.) at all time-intervals. (e.) The distribution of displacements between different molecules on the same ruler are well described by a model incorporating the localization precision (10nm) and the separation distance (50nm).

Simulated localizations were subjected to a spatiotemporal auto-correlation analysis as described in Methods and representative plots of the spatial component of *g*(*r,τ*) are shown in Fig 1b. This family of curves contains two major features: an initial peak at short displacements (r<40nm) arising from multiple localizations from the same molecule, and a second feature at wider radii (40nm<r<100nm) arising from displacements between localizations from different molecules on the same ruler. The amplitude of the initial peak decreases with increasing τ, while the second feature is largely independent of τ. The τ dependent component is isolated by subtracting *g*(*r, τ*) at long τ from those arising from shorterτ to obtain Δ*g*(*r, τ*) as shown in Fig 1d. In this simulation, there are no τ dependent effects that would impact resolution, resulting in Δ*g*(*r, τ*) having the same width for all τ. This is summarized by fitting Δ*g*(*r, τ*) to the Gaussian function of Eqn. 6 to extract the estimated width of the effective PSF *σ_xy_* which is reported in Fig 1e. In this simulated example, we can associate all localizations with the molecules that produced them, and can therefore directly compute *g_PSF_*(*r, τ*)from localizations as described in Methods and the resulting resolution obtained is also plotted in Fig 1e. Lastly, we tabulate displacements between all localizations originating from distinct molecules on the same ruler. The distribution is shown in Fig 1(f) and its properties are described by simulation parameters. The line in Fig 1f has a Gaussian shape with the form: 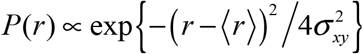 where *σ_xy_* is 10nm and 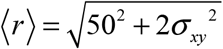 is 51.96nm. The slight bias in 〈*r*〉 towards a value larger than separation between molecules (50nm) arises from the components of localizations that fall perpendicular to the ruler axis and always contribute positive values to the measured displacements. For comparison, the FRC was tabulated to be 30nm for this simulation following procedures described in Methods. This FRC value is slightly larger than 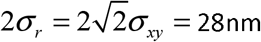.

The simulation of Fig 1 does not contain any factors expected to degrade image resolution over time. In Figure 2, the same simulation is subjected to a directed drift in the x-direction as well as diffusive drift in both x and y. Drift is corrected using a mean shift algorithm (20) that works by evenly dividing localizations into time-bins, then finding the displacement that minimizes the mean distance between localizations across all time-bins. The applied drift and the calculated drift correction are shown in Fig 2a along with the resulting image reconstruction, *g*(*r, τ*) curves over different windows in time-interval (Fig 2b.) closely resemble those shown in the static simulation, but Δ*g*(*r, τ*) now broadens with increasing τ (Fig 2c.). This broadening reflects a degradation of image resolution beyond the localization precision at all but the shortest time-intervals that plateaus beyond the timescale of the drift-correction (Fig 2d.). Here, the measured resolution reaches a local maximum at a time separation somewhat smaller than the drift correction timescale, which we attribute to the drift correction algorithm itself as it is also apparent in the resolution obtained by associating localizations with their originating molecules, as described in Methods. Deviations from the expected distribution of pairwise distances between localizations of molecules from opposite ends of the same nanoruler also exceed those of the static case (Fig 2e.) and are better described by a model that incorporates the measured image resolution, 〈*σ_xy_*〉 = 11.8*nm*, which is determined by averaging over estimated *σ_y_*(*τ*) weighted by the number of pairs associated with each time-interval window. The FRC resolution of the drift-corrected dataset in Figure 2 is 35 nm. This is somewhat larger than 2〈*σ_r_*〉 = 33*nm*. Observing a plateau in plots of *σ_xy_*(*τ*) is a good indicator that the algorithm is successfully applied, since drift correction is designed to stabilize errors on long time-scales. Supplementary Figure S3 shows an example of the same simulation with drift and drift correction, but where individual molecules are allowed to diffuse slowly such that *g_PSF_*(*r*) increases with τ in a way that is not accounted for through drift correction. In that case, *σ_xy_*(*τ*) increases with τ and is underestimated by Δ*g*(*r, τ*).

**Figure 2:**
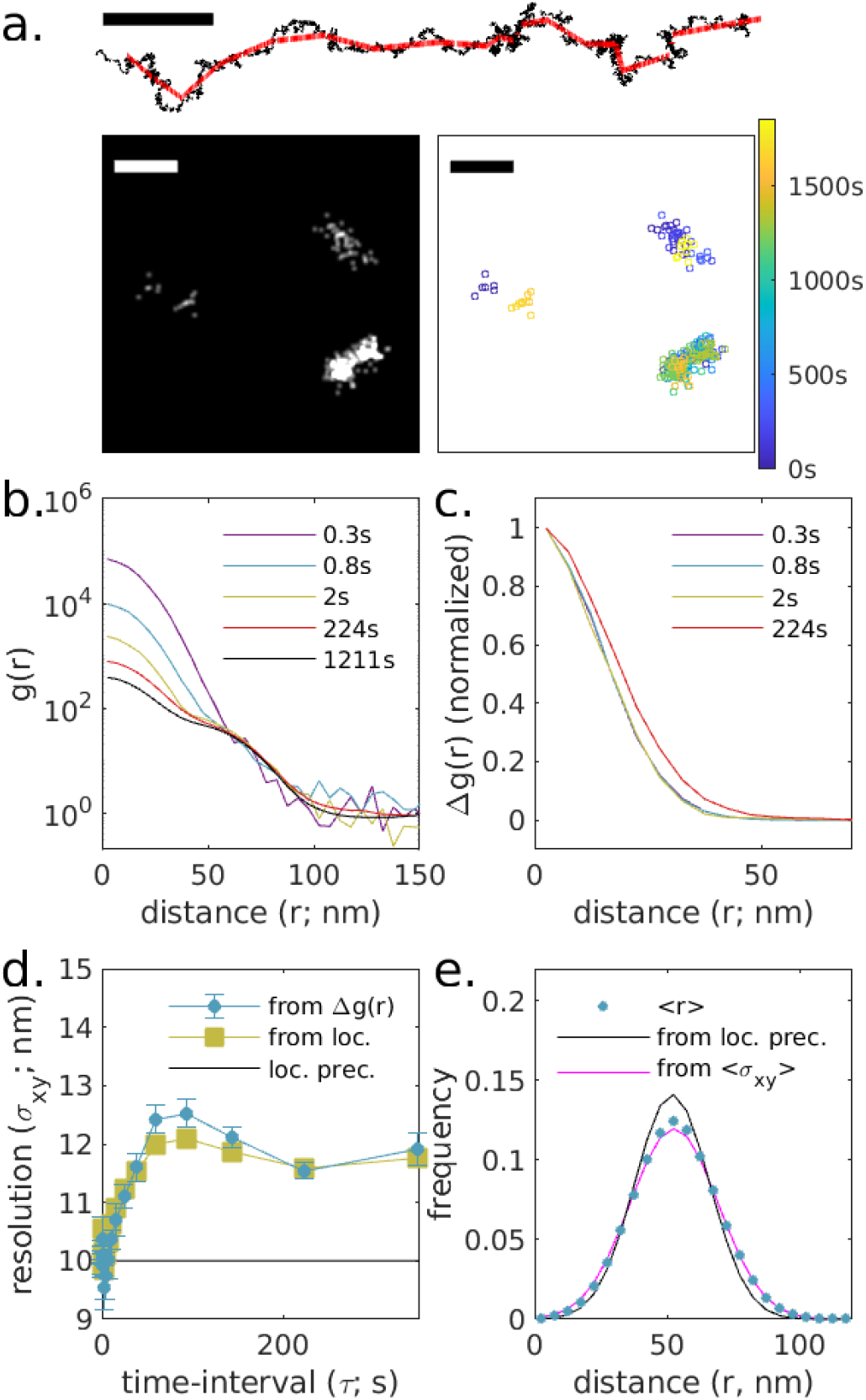
Validation of approach through simulation with drift and drift correction. (a.) The simulation from Fig 1 with applied drift (black) and drift correction (red) as shown in the trajectory above. Reconstructed image (left) and scatterplot of localizations with color representing the observation time (right) for a small subset of the simulated plane. Scale bar is 100nm. (b.) Auto-correlations as a function of displacement, *g*(*r, τ*), tabulated from simulations for time-interval windows centered at the values shown, (c.) Δ*g*(*r,τ*) = *g*(*r, τ*) – *g*(*r, τ*_max_ = 1211*s*) for the examples shown in b. (d.) Δ*g*(*r, τ*) are fit to 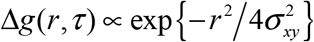 to extract the width of the effective point spread function in each lateral dimension. In this case, the resulting resolution estimate varies with time-interval, closely following the PSF measured by grouping localizations with molecules (from loc.). (e.) The distribution of displacements between different molecules on the same ruler are well described by a model incorporating the average estimated PSF (〈*σ_xy_*〉 = 118*nm*) and the separation distance (50nm).

### Estimating the effective PSF of experimental SMLM datasets

Figure 3 demonstrates this approach on an experimental dataset of DNA origami nanorulers that resemble the simulated rulers with AlexaFluor647 labeling sites separated by 50nm. Fig 3a shows a small subset of the field of view of the acquired image, that was reconstructed from 29,000 image frames acquired over 53 min at a frame rate of 0.1s, with a total of over 126,000 individual localizations. In post-processing, a drift correction was applied with a time-window width of 25s or 250 image frames. As in the simulated case, *g*(*r, τ*) decays at short r with increasing *τ* (Fig 3c), and Δ*g*(*r, τ*) is roughly Gaussian (Fig 3d). Fitting Δ*g*(*r, τ*) yields the resolution *σ_xy_*(*τ*). As in the simulated example, the measured resolution is lowest at short time-intervals (close to 7nm) and plateaus at time-scales somewhat shorter than the frequency of the applied drift correction.

**Figure 3:**
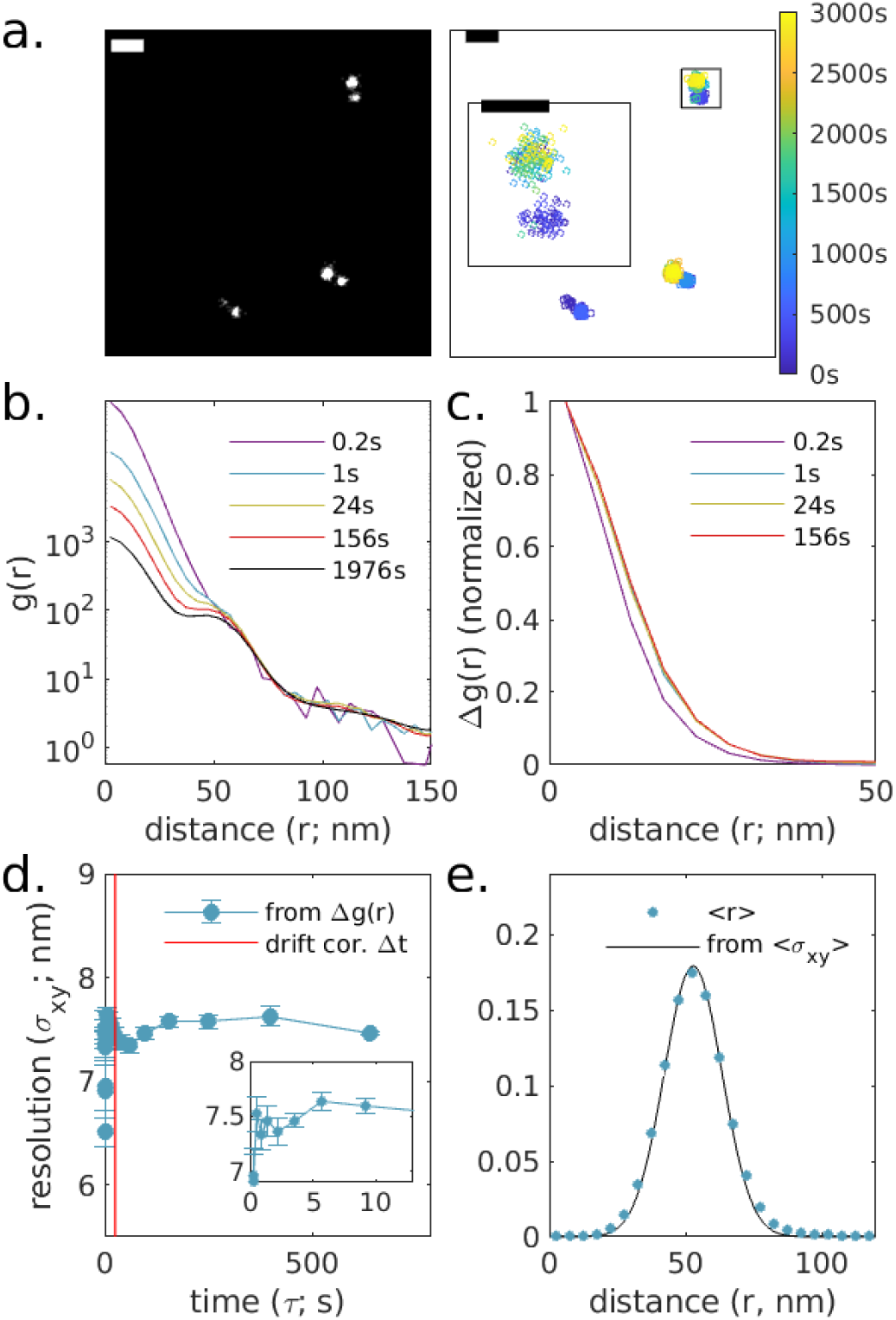
Experimental observations of DNA origami rulers labeled with AlexaFluor647. (a.) Reconstructed image (left) and scatterplot of localizations with color representing the observation time (right) for a small subset of the observed plane. Scale bar is 100nm and 50nm in the inset. A larger field of view from this image is shown in Supplementary Figure S5. (b.) Auto-correlations as a function of displacement, *g*(*r, τ*), tabulated from localizations for time-interval windows centered at the values shown. (c.) Δ*g*(*r, τ*) = *g*(*r, τ*) – *g*(*r, τ*_max_ = 1976*s*)for the examples shown in b. (d.) Δ*g*(*r, τ*) are fit to 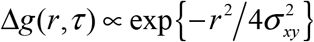 to extract the width of the effective point spread function in each lateral dimension which varies with time-interval. The resulting average resolution for this image is 〈*σ_xy_*〉 = 7.4*nm*. (e.) The distribution of displacements between pairs of fluorophores on the same ruler, segmented using DBSCAN as described in Methods. Fitting to a Gaussian shape with width given by the measured 〈*σ_xy_*〉 produces 〈*r*〉 = 52.2±0.2*nm*.

Since the localization clouds from individual Alexa647 molecules were visually distinct, we applied a DBSCAN segmentation algorithm (29) to associate localizations with individual molecules. From this segmentation, we tabulated the pairwise distances between molecules on the same origami and the distribution of these values is shown in Fig 3e. This distribution is well described by a model applying the measured 〈*σ_xy_*(*τ*)〉 = 7.5*nm* with 〈*r*〉 = 52.2± 0.2 nm, where the error is dominated by uncertainty in the sample magnification at the camera. This yields a separation distance of 51.1 ± 0.2*nm* between labels on individual rulers, which is within the manufacturer’s specifications. The FRC resolution was found to be 19 nm, slightly smaller than 2〈*σ_r_*〉 = 21.2*nm*.

We have conducted this same analysis on a similar DNA origami sample that was imaged using DNA PAINT, this time using rulers containing 3 collinear docking sites separated by 80nm and summarized in Figure 4. In contrast to the dSTORM fluorophores of Fig 3, molecules imaged by DNA PAINT do not exhibit long time-scale correlations, limiting the applicability of this method. The DNA PAINT probes used for this image do remain correlated over time-scales relevant for drift-correction (~15s), which is long enough to provide a useful estimate of image resolution. In this example, drift correction was applied with a time-window width of 11s or 110 image frames. Pairwise distances between labels on the center and ends of the origami were measured after applying DBSCAN to segment localizations from distinct docking sites and are well described by a model applying the measured 〈*σ_xy_*(*τ*)〉 = 8.7*nm* with 〈*r*} = 82.5± 0.3*nm*. This yields a separation distance of 81.6±0.3*nm* between the center and endpoint labels on individual rulers. Since temporal correlations of the PAINT probes used in this example only extend for a small fraction of the acquisition time (20 min), the average *σ_xy_* is given primarily by the value determined in the largest time-interval bin (*τ* = 15*s*). This is because the vast majority of pairs are detected at time-intervals grouped into *τ*_max_, where we do not estimate *σ_xy_* but instead apply the value estimated at the previous time-window bin. The good agreement between the model and measured distributions in Fig. 3e validates this approach, at least for this specific example where drift correction was accomplished on a shorter time-scale. The FRC resolution was found to be 26 nm, slightly larger than 2〈*σ_r_*〉 = 24.6*nm*.

**Figure 4:**
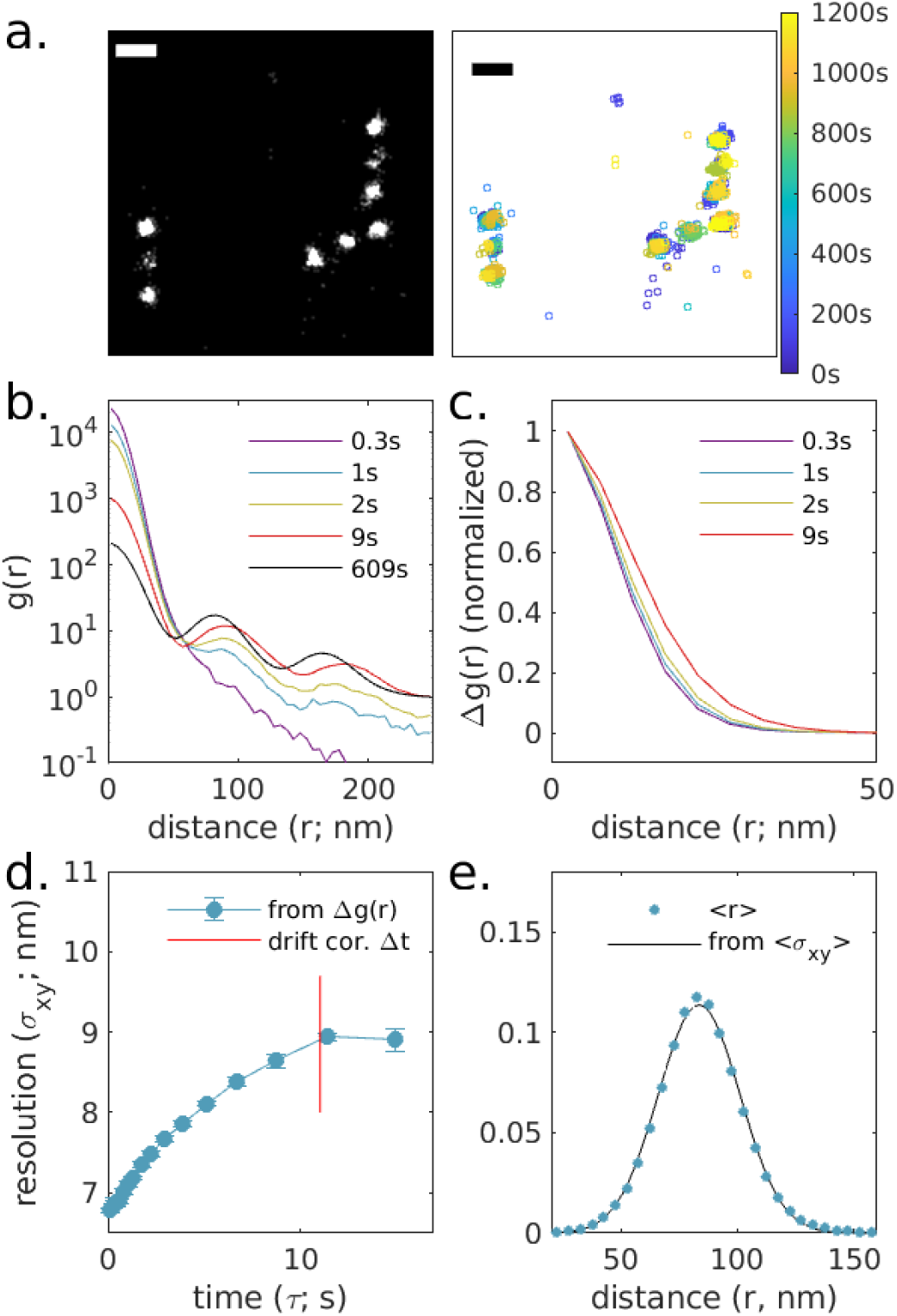
Experimental observations of DNA origami rulers imaged with DNA PAINT, using an Atto655 imaging strand, (a.) Reconstructed image (left) and scatterplot of localizations with color representing the observation time (right) for a small subset of the observed plane. Scale bar is lOOnm and 5Onm in the inset. A larger field of view from this image is shown in Supplementary Figure S6. (b.) Auto-correlations as a function of displacement, *g*(*r, τ*), tabulated from localizations for time-interval windows centered at the values shown, (c.) Δ*g*(*r, τ*) = *g*(*r, τ*) – *g*(*r,τ*_max_ = 609*s*) for the examples shown in b. (d.) Δ*g*(*r, τ*) are fit to 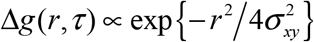 to extract the width of the effective point spread function in each lateral dimension which varies with time-interval. The resulting average resolution for this image is 〈*σ_xy_*〉 = 8.7*nm*. (e.) The distribution of displacements between different molecules on the same ruler, segmented using DBSCAN as described in Methods. Fitting to a Gaussian shape with width given by the measured 〈*σ_xy_*〉 produces 〈*r*〉 = 82.5±0.3*nm*.

We next apply this method to image labeled structures in cells. Figure 5 shows the method applied to nuclear pore complexes (NPCs) within the nuclear envelope of chemically fixed primary mouse neurons. In these images, a protein component of NPCs, NUP210, was labeled with a conventional primary antibody and a Fab secondary directly conjugated to AlexaFluor647. 12500 images were acquired over 23 min with an integration time of 0.1s and a total of 178873 localizations detected within the masked ROI at the nuclear envelope. Drift correction was accomplished with a time-window of 8.3 s or 83 image frames. Reconstructed images of the entire nucleus and single pores are shown in Fig 5a. along with a scatter plot demonstrating that individual NPC subunits are sampled at times throughout the observation, *g*(*r, τ*) (Fig 5b.) curves extend to beyond 100nm reflecting the extended structure of individual labeled NPCs, but extended structure is effectivly removed by examining Δ*g*(*r,τ*) (Fig 5c.). Fitting Δ*g*(*r, τ*) to a Gaussian shape quantifies image resolution over time, which is larger than the localization precision beyond the shortest time-interval shown. We estimate 〈*σ_xy_*〉 to be 10.9nm. The FRC resolution was evaluated to be 37 nm, which is larger than 2〈*σ_r_*〉 = 30.8*nm*.

**Figure 5:**
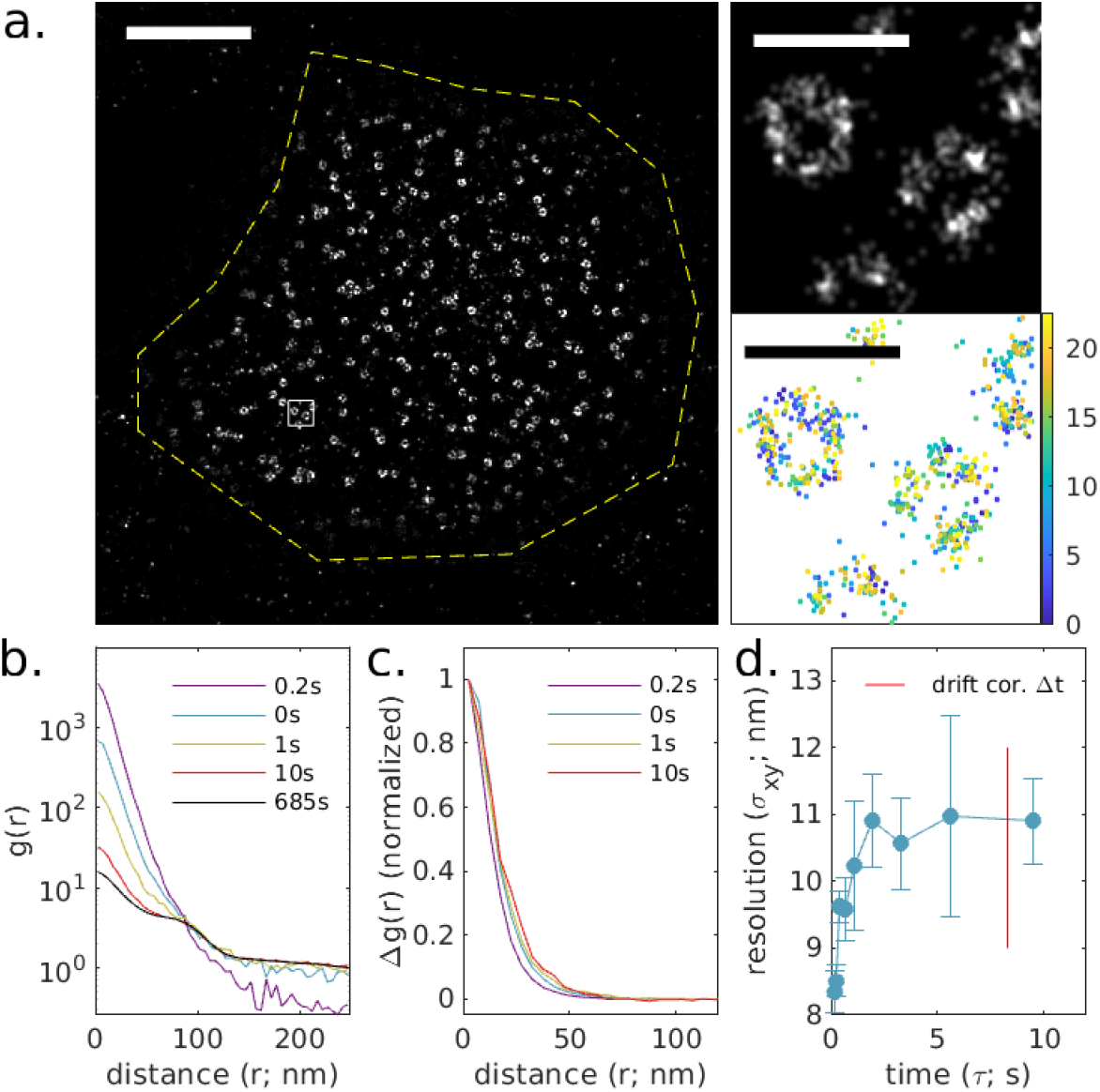
Experimental observations of Nuclear Pore Complexes (NPCs) within primary mouse neurons, antibody-labeled with AlexaFluor647. (a.) Reconstructed image (left) and image showing region of interest interrogated (yellow dashed region) and a magnified subset (top right, from white square region of larger image) along with a scatterplot of localizations with color representing the observation time (bottom right). Scale-bar is 2μm (left) and 200nm (right top and bottom), (b.) Auto-correlations as a function of displacement, *g*(*r, τ*), tabulated from localizations for time-interval windows centered at the values shown, (c.) Δ*g*(*r, τ*) = *g*(*r, τ*) – *g*(*r, τ*_max_ = 685*s*)for the examples shown in b. (d.) Δ*g*(*r, τ*) are fit to 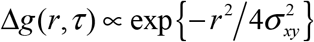 to extract the width of the effective point spread function in each lateral dimension. The resulting average resolution for this image is 〈*σ_xy_*〉 = 10.9*nm*.

Figure 6 shows a similar class of cellular structure imaged using DNA PAINT. In this example, clathrin-gfp is transiently expressed in CH27 cells then labeled post fixation with an anti GFP nanobody conjugated to an ssDNA docking strand. Cells are then imaged in the presence of a complementary imaging strand labeled with Atto 655. Similar to the origami DNA PAINT sample of Figure 4, temporal correlations from single molecules remain for short to medium time-scales (~8sec), allowing for accurate estimation of resolution losses due to drift and drift correction. Here we estimate the resolution 〈*σ_xy_*〉 to be 11.6nm. The FRC resolution was evaluated to be 47 nm, which is larger than 2〈*σ_r_*〉 = 32.8*nm*.

**Figure 6:**
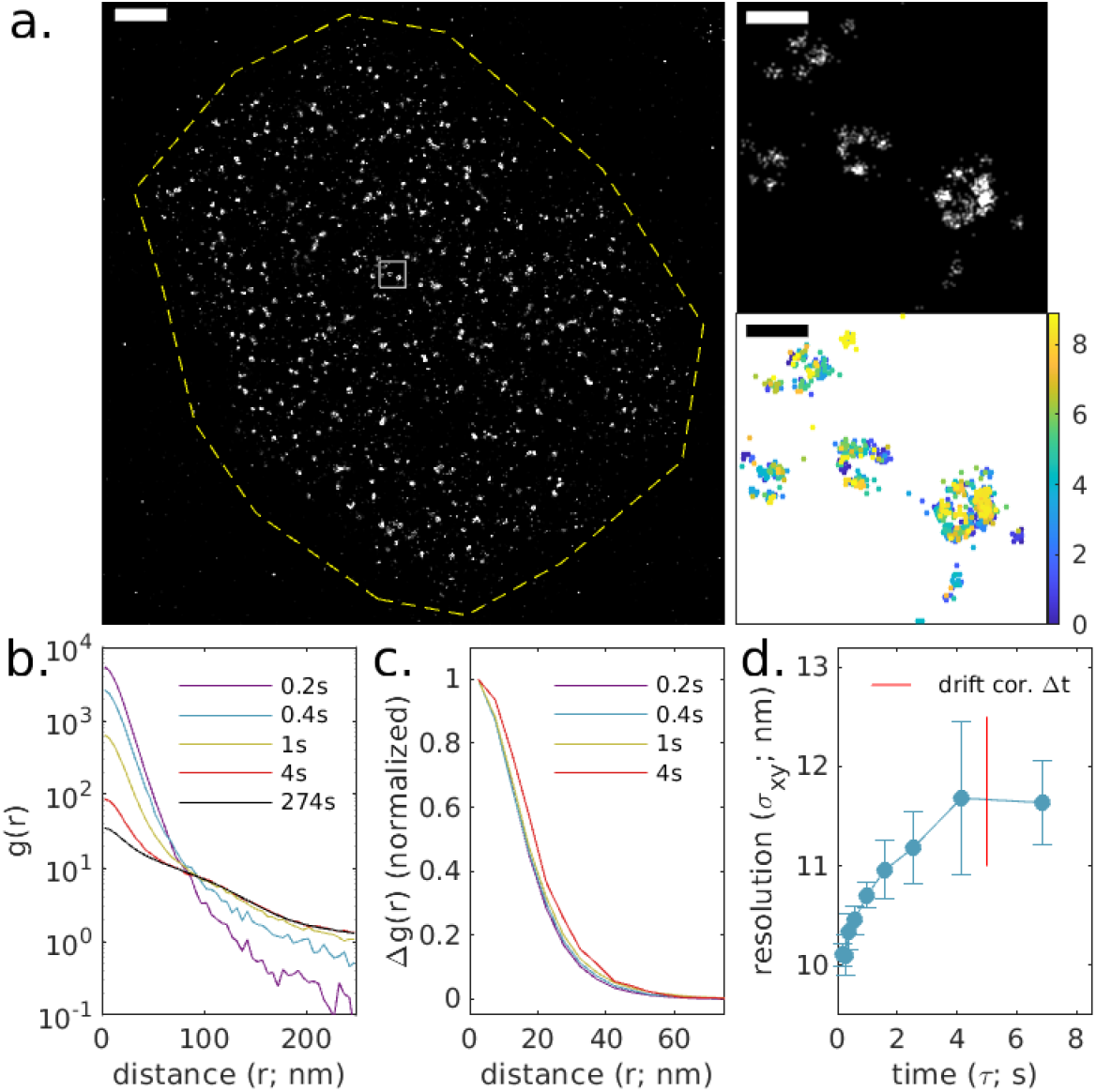
Experimental observations of clathrin coated pits within CH27 B cells, imaged using a nanobody-coupled Atto655 DNA-PAINT scheme. (a.) Reconstructed image (left) and image showing region of interest interrogated (yellow dashed region) and a magnified subset (top right, from white square region of larger image) along with a scatterplot of localizations with color representing the observation time (bottom right). Scale-bar is 2μm (left) and 200nm (right top and bottom), (b.) Auto-correlations as a function of displacement, *g*(*r, τ*), tabulated from localizations for time-interval windows centered at the values shown. (c.) Δ*g*(*r, τ*) = *g*(*r, τ*) – *g*(*r, τ*_max_ = 274*s*) for the examples shown in b. (d.) Δ*g*(*r, τ*) are fit to 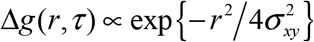 to extract the width of the effective point spread function in each lateral dimension. The resulting average resolution for this image is 〈*σ_xy_*〉 = 11.6*nm*.

Figure 7 shows the method applied to an image of f-actin staining by phalloidin-AlexaFluor647 in chemically fixed CH27 B cells adhered to a glass surface decorated with VCAM. For this sample, 5000 images were acquired over 4.9 min with an integration time of 0.05s and a total of 302681 localizations within the masked ROI. Drift correction was accomplished with a time-window of 2.5s or 50 image frames. Unlike Figs 5 and 6 where labels decorate isolated structures scattered over a surface, this reconstructed image of f-actin is more space filling, making up a web of fibers that extend across the entire ventral cell surface (Fig 7a.). This extended structure can be detected in *g*(*r, τ*) (Fig 7b.) as increased intensity in the tail that extends to large separation distances for curves generated at all τ. This large-scale structure is effectively removed in Δ*g*(*r,τ*) (Fig 7c) allowing for a determination of the effective PSF over a range of time-scales as shown in Fig 7d. In this example, the ROI was drawn within the cell boundary to minimize the intensity of *g_p_*(*r*) which allows for accurate estimation of *g_PSF_*(*r,τ*) out to longer time-intervals. This is because the amplitude of *g*(*r, τ*_max_) includes contributions from *g_p_*(*r*), while the amplitude of Δ*g*(*r, τ*) only depends on *g_e_*(*τ*), therefore Δ*g*(*r* < 25*nm,τ*)/*g*(*r* < 25*nm, τ*_max_) will remain larger than the cutoff over a wider range of *τ*. The estimate for 〈*τ_xy_*〉 is 11.8nm. The FRC resolution was found to be 290 nm, which is much larger than 2〈*σ_r_*〉 = 33*nm*. This large discrepancy likely reflects the under-sampling of actin filaments in this image.

**Figure 7:**
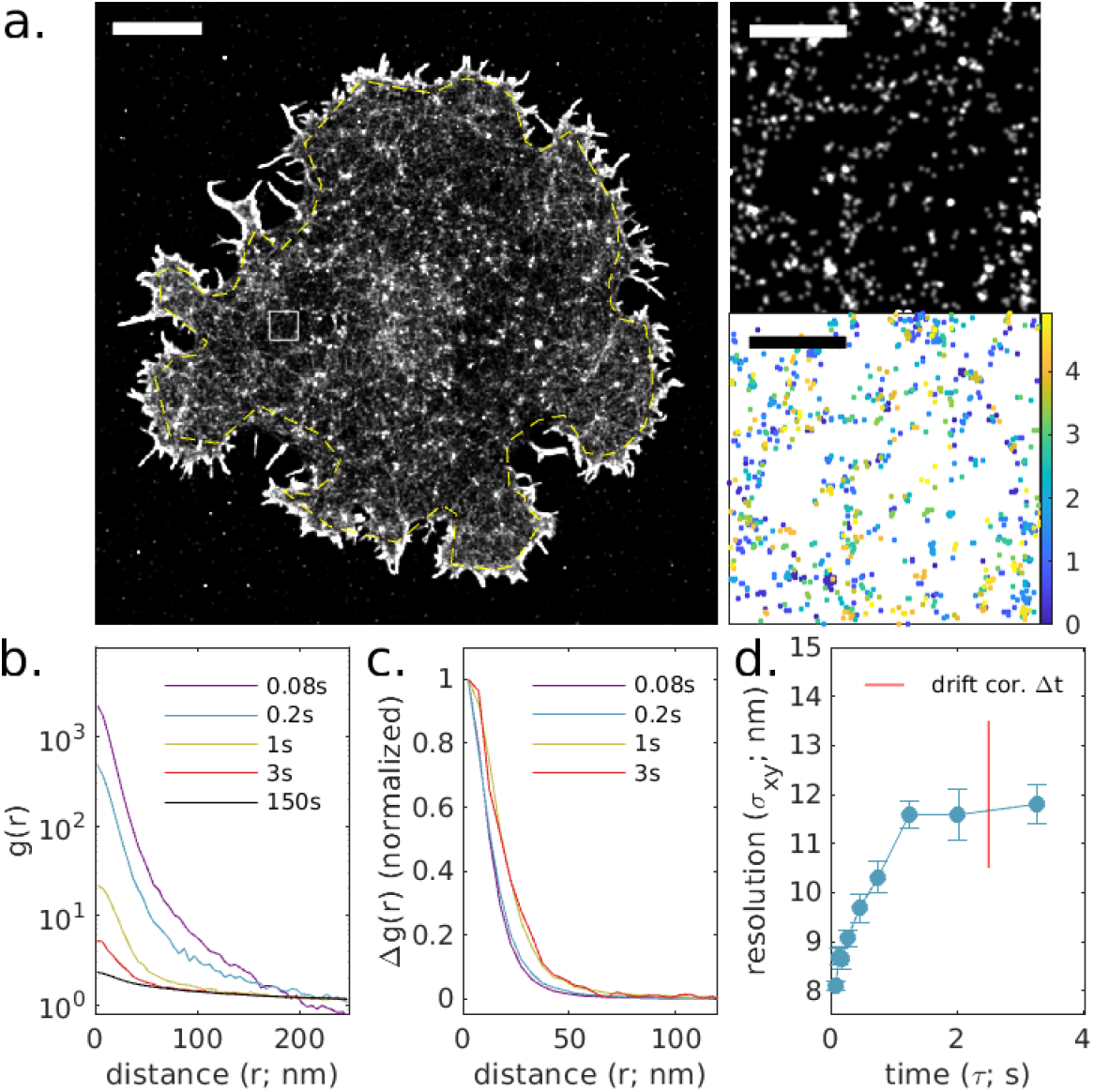
Experimental observations of f-actin on the ventral surface of a CH27 B cell using phalloidin-AlexaFluor647. (a.) Reconstructed image (left) and image showing region of interest interrogated (yellow dashed region inside cell boundary) and a magnified subset (top right, from white square region of larger image) along with a scatterplot of localizations with color representing the observation time (bottom right). Scale-bar is 5μm (left) and 500nm (right top and bottom). (b.) Auto-correlations as a function of displacement, *g*(*r, τ*), tabulated from localizations for timeinterval windows centered at the values shown. (c.) Δ*g*(*r, τ*) = *g*(*r, τ*) – *g*(*r, τ*_max_ = 150*s*)for the examples shown in b. (d.) Δ*g* (*r, τ*) are fit to 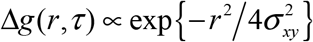 to extract width of the effective point spread function in each lateral dimension. The time-scale of the drift correction is shown in red. The resulting average resolution for this image is 〈*σ_xy_*〉 = 11.8*nm*.

As a final demonstration, Figure 8 shows the method applied to an image of Src15-mEos3.2, a myristoylated peptide bound to the inner leaflet of the plasma membrane and directly conjugated to the photo-switchable protein fluorophore mEos3.2. This peptide uniformly decorates the ventral surface of a chemically fixed CH27 B cell adhered to a glass surface decorated with VCAM, as seen in the reconstructed image of Fig 8a. For this sample, 7000 images were acquired over 12.7 min with an integration time of 0.1s and a total of 240,503 localizations. Drift correction was accomplished with a time-window of 12.5s or 125 image frames. mEos3.2 exhibits different blinking dynamics than AlexaFluor647, with some probes exhibiting correlated blinking on long time-scales. This can be seen in plots of *g*(*r, τ*) that take long timescales to decay (Fig 8b.). Again, Δ*g*(*r, τ*) curves isolate the initial peak, allowing for the quantification of the effective PSF. In this example, the slow decay of *g*(*r* < 25*nm, τ*) with τ allows for estimation of *σ_xy_* out to large time-intervals. The FRC resolution was found to be 57 nm, which is larger than 2〈*σ_r_*〉 = 39 nm.

**Figure 8:**
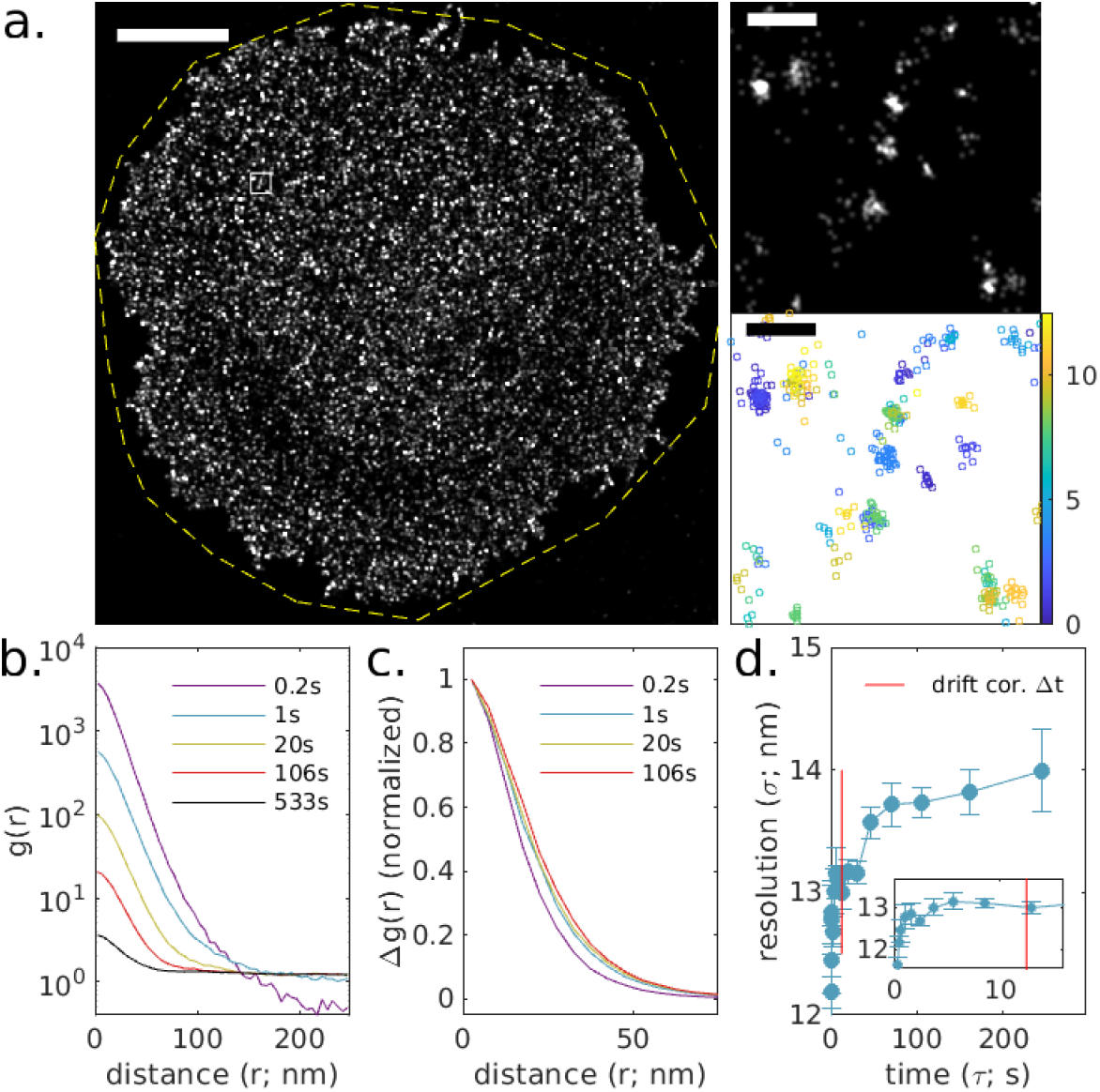
Experimental observations of membrane anchor peptide Src15-mEos3.2 on the ventral surface of a CH27 B cell. (a.) Reconstructed image (left) and image showing region of interest interrogated (yellow dashed region) and a magnified subset (top right, from white square region of larger image) along with a scatterplot of localizations with color representing the observation time (bottom right). Scale bar is 5μm (left) and 200nm (right top and bottom), (b.) Auto-correlations as a function of displacement, *g*(*r, τ*), tabulated from localizations for time-interval windows centered at the values shown, (c.) Δ*g*(*r, τ*) = *g*(*r, τ*) – *g*(*r, τ*_max_ = 533*s*) for the examples shown in b. (d.) Δ*g* (*r, τ*) are fit to 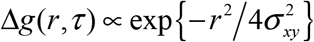 to extract the width of the effective point spread function in each lateral dimension. The time-scale of the drift correction is shown in red. The resulting average resolution for this image is 〈*σ_xy_*〉 = 13.7*nm*.

## Conclusions

Here we present a method to estimate the effective PSF of a super-resolution fluorescence localization measurement directly from acquired localizatoins, relying on a few reasonable assumptions. The basic method is validated through simulations and demonstrated using experimental data of three commonly used localization microscopy probes. Importantly, the described method performs best when used alongside flourophores exhibiting blinking dynamics that remain correlated in time out to time-scales relevant to sources of error present in the imaging experiment. This resolution metric directly reports on how accurately the positions of molecules are recorded at the end of an experimental and analytical pipeline, allowing experimenters to validate and optimize new methods. Furthermore, it explicitly probes time-dependent errors that emerge due to motion of the microscope stage and/or individual fluorophores. Beyond optimization, we expect this method to be useful when interpreting experiments that involve the measurement of distances between localizations in images, for example in auto- or cross-correlation analysis (6, 32, 33). It is a complementary approach to resolution metrics that also incorporate information about spatial sampling, most notably the FRC (12, 13).

The reported method estimates resolution by fitting the estimated effective PSF at each time-interval probed to a Gaussian shape, followed by a weighted average to extract the estimated error of the average localization in an image. This approach is convenient because the resolution is summarized as a single number. However, it may be possible to extract more detailed information about the PSF by loosening the assumption of a Gaussian shape. We anticipate that the full average effective PSF could prove useful for other purposes, such as deconvolution of reconstructed images or tabulated spatial correlation functions, or as input to clustering algorithms or other analysis tools.

Finally, the values obtained using the described method depend on long time-scale correlations of fluorescent probes used for imaging. For the examples shown, we find that it is most important to characterize resolution lost on time-scales shorter than the time-scale of drift-correction. Fortunately, numerous methods exist to correct for rigid drift on time-scales relevant to the temporal correlations of many SMLM probes(34–41), suggesting that the method presented in this report is broadly applicable for a range of experimental conditions.

## AUTHOR CONTRIBUTIONS

TRS, FJF and SLV developed the method. TRS and SLV developed the analysis code and performed the analyses, with help from FJF. TRS, FJF, and SLV developed the simulated data. SLV imaged the nano-ruler samples, and the clathrin, actin, and Src15, samples. The Src15 and clathrin samples were prepared by ERS and the actin sample was prepared by JCF-N. NPC sample preparation and imaging was carried out by SK. SLV and TRS wrote the text in consultation with the other authors.

## ACKNOWLEDGEMENTS

We thank Kathleen Wisser, Andrea Stoddard and Patrick DeLear for assistance with experiments. This work was supported by grants from the U.S. National Science Foundation (MCB1552439) and National Institutes of Health (GM129347 and GM110052).

## CITED REFERENCES

1. Rust, M.J., M. Bates, and X. Zhuang. 2006. Sub-diffraction-limit imaging by stochastic optical reconstruction microscopy (STORM). Nat. Methods. 3:793–796.

2. Heilemann, M., E. Margeat, R. Kasper, M. Sauer, and P. Tinnefeld. 2005. Carbocyanine Dyes as Efficient Reversible Single-Molecule Optical Switch. J. Am. Chem. Soc. 127:3801–3806.

3. Hess, S.T., T.P.K. Girirajan, and M.D. Mason. 2006. Ultra-High Resolution Imaging by Fluorescence Photoactivation Localization Microscopy. Biophys. J. 91:4258–4272.

4. Betzig, E., G.H. Patterson, R. Sougrat, O.W. Lindwasser, S. Olenych, J.S. Bonifacino, M.W. Davidson, J. Lippincott-Schwartz, and H.F. Hess. 2006. Imaging Intracellular Fluorescent Proteins at Nanometer Resolution. Science. 313:1642–1645.

5. Sharonov, A., and R.M. Hochstrasser. 2006. Wide-field subdiffraction imaging by accumulated binding of diffusing probes. Proc. Natl. Acad. Sci. 103:18911–18916.

6. Veatch, S.L., B.B. Machta, S.A. Shelby, E.N. Chiang, D.A. Holowka, and B.A. Baird. 2012. Correlation Functions Quantify Super-Resolution Images and Estimate Apparent Clustering Due to Over-Counting. PLoS ONE. 7:e31457.

7. Endesfelder, U., S. Malkusch, F. Fricke, and M. Heilemann. 2014. A simple method to estimate the average localization precision of a single-molecule localization microscopy experiment. Histochem. Cell Biol. 141:629–638.

8. Demmerle, J., E. Wegel, L. Schermelleh, and I.M. Dobbie. 2015. Assessing resolution in super-resolution imaging. Methods. 88:3–10.

9. Descloux, A., K.S. Grußmayer, and A. Radenovic. 2019. Parameter-free image resolution estimation based on decorrelation analysis. Nat. Methods. 16:918–924.

10. Saxton, W.O., and W. Baumeister. 1982. The correlation averaging of a regularly arranged bacterial cell envelope protein. J. Microsc. 127:127–138.

11. Van Heel, M. 1987. Similarity measures between images. Ultramicroscopy. 21:95–100.

12. Nieuwenhuizen, R.P.J., K.A. Lidke, M. Bates, D.L. Puig, D. Grünwald, S. Stallinga, and B. Rieger. 2013. Measuring image resolution in optical nanoscopy. Nat. Methods. 10:557–562.

13. Banterle, N., K.H. Bui, E.A. Lemke, and M. Beck. 2013. Fourier ring correlation as a resolution criterion for super-resolution microscopy. J. Struct. Biol. 183:363–367.

14. Annibale, P., S. Vanni, M. Scarselli, U. Rothlisberger, and A. Radenovic. 2011. Quantitative Photo Activated Localization Microscopy: Unraveling the Effects of Photoblinking. PLOS ONE. 6:e22678.

15. Nieuwenhuizen, R.P.J., M. Bates, A. Szymborska, K.A. Lidke, B. Rieger, and S. Stallinga. 2015. Quantitative Localization Microscopy: Effects of Photophysics and Labeling Stoichiometry. PLOS ONE. 10:e0127989.

16. Patel, L., D. Williamson, D.M. Owen, and E.A.K. Cohen. 2021. Blinking statistics and molecular counting in direct stochastic reconstruction microscopy (dSTORM). Bioinformatics. 37:2730–2737.

17. Raab, M., I. Jusuk, J. Molle, E. Buhr, B. Bodermann, D. Bergmann, H. Bosse, and P. Tinnefeld. 2018. Using DNA origami nanorulers as traceable distance measurement standards and nanoscopic benchmark structures. Sci. Rep. 8:1780.

18. Patel, L., N. Gustafsson, Y. Lin, R. Ober, R. Henriques, and E. Cohen. 2019. A hidden Markov model approach to characterizing the photo-switching behavior of fluorophores. Ann. Appl. Stat. 13:1397–1429.

19. Dana. 2022. Simulate Continuous-Time Markov Chains.

20. Fazekas, F.J., T.R. Shaw, S. Kim, R.A. Bogucki, and S.L. Veatch. 2021. A mean shift algorithm for drift correction in localization microscopy. Biophys. Rep. 1:100008.

21. Haughton, G., L.W. Arnold, G.A. Bishop, and T.J. Mercolino. 1986. The CH Series of Murine B Cell Lymphomas: Neoplastic Analogues of Ly-1+ Normal B Cells. Immunol. Rev. 93:35–52.

22. Stone, M.B., and S.L. Veatch. 2015. Steady-state cross-correlations for live two-colour super-resolution localization data sets. Nat. Commun. 6.

23. Chan, J.R., S.J. Hyduk, and M.I. Cybulsky. 2000. α4β1 Integrin/VCAM-1 Interaction Activates αLβ2 Integrin-Mediated Adhesion to ICAM-1 in Human T Cells. J. Immunol. 164:746–753.

24. Rodgers, W. 2002. Making membranes green: construction and characterization of GFP-fusion proteins targeted to discrete plasma membrane domains. BioTechniques. 32:1044–1046, 1048, 1050–1051.

25. Zhang, M., H. Chang, Y. Zhang, J. Yu, L. Wu, W. Ji, J. Chen, B. Liu, J. Lu, Y. Liu, J. Zhang, P. Xu, and T. Xu. 2012. Rational design of true monomeric and bright photoactivatable fluorescent proteins. Nat. Methods. 9:727–729.

26. Izeddin, I., J. Boulanger, V. Racine, C.G. Specht, A. Kechkar, D. Nair, A. Triller, D. Choquet, M. Dahan, and J.B. Sibarita. 2012. Wavelet analysis for single molecule localization microscopy. Opt. Express. 20:2081–2095.

27. Smith, C.S., N. Joseph, B. Rieger, and K.A. Lidke. 2010. Fast, single-molecule localization that achieves theoretically minimum uncertainty. Nat. Methods. 7:373–375.

28. Ovesný, M., P. Křížek, J. Borkovec, Z. Švindrych, and G.M. Hagen. 2014. ThunderSTORM: a comprehensive ImageJ plug-in for PALM and STORM data analysis and super-resolution imaging. Bioinformatics. 30:2389–2390.

29. Ester, M., H.-P. Kriegel, J. Sander, and X. Xu. 1996. A density-based algorithm for discovering clusters in large spatial databases with noise. AAAI Press. pp. 226–231.

30. Dai, M., R. Jungmann, and P. Yin. 2016. Optical imaging of individual biomolecules in densely packed clusters. Nat. Nanotechnol. 11:798–807.

31. Shaw, T.R., F.J. Fazekas, and S.L. Veatch. 2022. VeatchLab / SMLM Spacetime Resolution. https://gitlab.umich.edu/veatchlab/smlm-spacetime-resolution.

32. Stone, M.B., S.A. Shelby, M.F. Núñez, K. Wisser, and S.L. Veatch. 2017. Protein sorting by lipid phase-like domains supports emergent signaling function in B lymphocyte plasma membranes. eLife. 6:e19891.

33. Andersen, I.T., U. Hahn, E.C. Arnspang, L.N. Nejsum, and E.B.V. Jensen. 2018. Double Cox cluster processes — with applications to photoactivated localization microscopy. Spat. Stat. 27:58–73.

34. Colomb, W., J. Czerski, J.D. Sau, and S.K. Sarkar. 2017. Estimation of microscope drift using fluorescent nanodiamonds as fiducial markers: ESTIMATION OF MICROSCOPE DRIFT. J. Microsc. 266:298–306.

35. Ma, H., J. Xu, J. Jin, Y. Huang, and Y. Liu. 2017. A Simple Marker-Assisted 3D Nanometer Drift Correction Method for Superresolution Microscopy. Biophys. J. 112:2196–2208.

36. Balinovic, A., D. Albrecht, and U. Endesfelder. 2019. Spectrally red-shifted fluorescent fiducial markers for optimal drift correction in localization microscopy. J. Phys. Appl. Phys. 52:204002.

37. Wang, Y., J. Schnitzbauer, Z. Hu, X. Li, Y. Cheng, Z.-L. Huang, and B. Huang. 2014. Localization events-based sample drift correction for localization microscopy with redundant cross-correlation algorithm. Opt. Express. 22:15982.

38. Elmokadem, A., and J. Yu. 2015. Optimal Drift Correction for Superresolution Localization Microscopy with Bayesian Inference. Biophys. J. 109:1772–1780.

39. Schlangen, I., J. Franco, J. Houssineau, W.T.E. Pitkeathly, D. Clark, I. Smal, and C. Rickman. 2016. Marker-Less Stage Drift Correction in Super-Resolution Microscopy Using the Single-Cluster PHD Filter. IEEE J. Sel. Top. Signal Process. 10:193–202.

40. Wester, M.J., D.J. Schodt, H. Mazloom-Farsibaf, M. Fazel, S. Pallikkuth, and K.A. Lidke. 2021. Robust, fiducial-free drift correction for super-resolution imaging. Sci. Rep. 11:23672.

41. Smith, C., chirlmin joo, jelmer Cnossen, and T.J. Cui. 2021. Drift correction in localization microscopy using entropy minimization. Opt. Express.

