## Supplementary figures and notes for "A method to estimate the effective point spread function of static single molecule localization microscopy images"

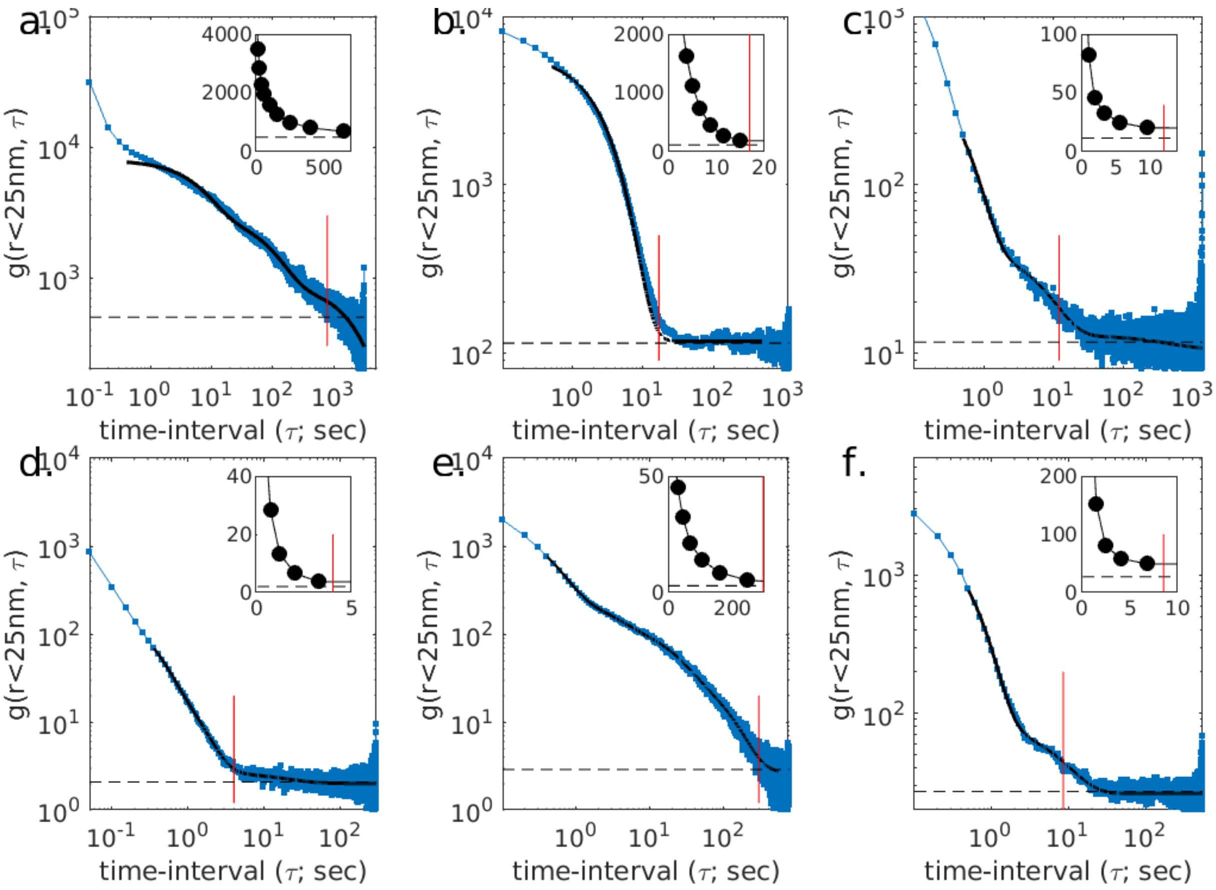


**Supplementary Figure 1.** Plots of $g(r < 25nm, \tau)$ for the experimental samples shown the main text: (a.) Alexa647 DNA origami rulers from Fig 3, (b.) PAINT DNA origami rulers from Fig 4, (c.) Alexa647 labeled NUP 210 from Fig 5, (d.) Alexa647-phalloidin from Fig 6. (e.) mEeos3.2 conjugated Src15 from Fig 7. (f.) PAINT labeled clathrin from Fig 8. These curves capture $g_e(\tau)$ up to a numerical offset that is dependent on the structure present in the image. Black lines are fit to a sum of exponentials and are present to highlight the monotonically decreasing trend. Dashed lines indicate the average value over the last ¼ of the dataset. The red vertical line indicates where $g(r < 25nm, \tau)$ falls below 1.5 times the dashed line, indicating the lower bin edge used to define $\tau_{\max}$.

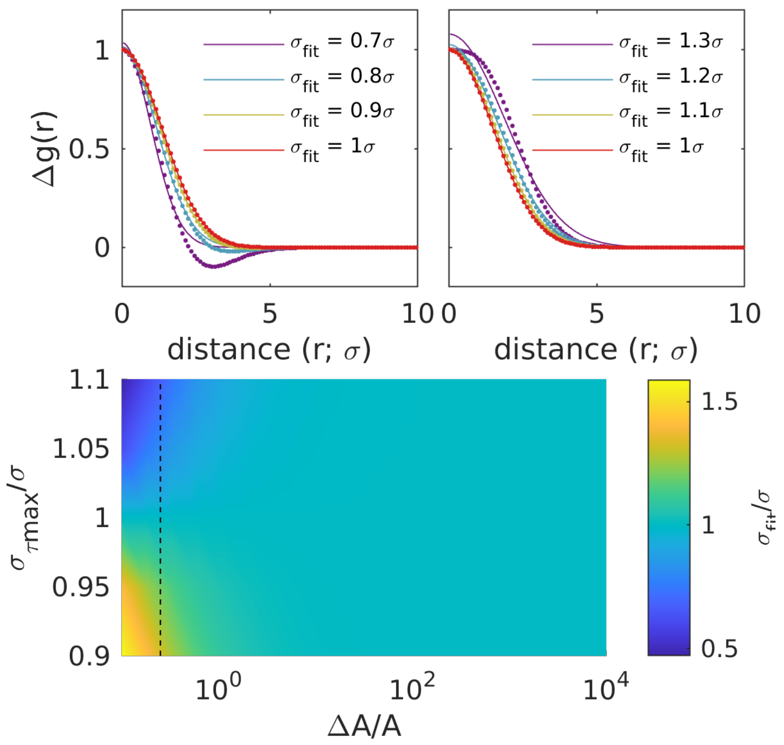


**Supplementary Figure S2:** Subtracting Gaussian shapes with different width leads to distortion in $\Delta g(r) = g(r,\tau) - g(r,\tau_{\max})$ when $g(r,\tau)$ and $g(r,\tau_{\max})$ have similar amplitudes but different widths. (top) plots of $\Delta g(r) = (A+\Delta A)\exp\left\{-r^2/4\sigma^2\right\} - A\exp\left\{-r^2/4\sigma_{\tau\max}{}^2\right\}$ for $\sigma_{\tau\max} = 1.1\sigma$ (left) and $\sigma_{\tau\max} = 0.9\sigma$ (right) and $\Delta A = 0.25, 0.5, 1, 2$ from purple to red. Curves are normalized so they pass through 1 at r=0. The legend shows the width extracted when fitting $\Delta g(r)$ to a single Gaussian shape $\Delta g(r) = A\exp\left\{-r^2/4\sigma_{fit}{}^2\right\}$. A broader $\sigma_{\tau\max}$ leads to systematic narrowing of $\sigma_{fit}$, while a narrow $\sigma_{\tau\max}$ leads to systematic broadening of $\sigma_{fit}$ when the difference in amplitudes is order 1. (bottom) a summary of results over a broad range of $\Delta A$ and $\sigma_{\tau\max}$ indicates that distortion is not a major concern over broad range of values interrogated.

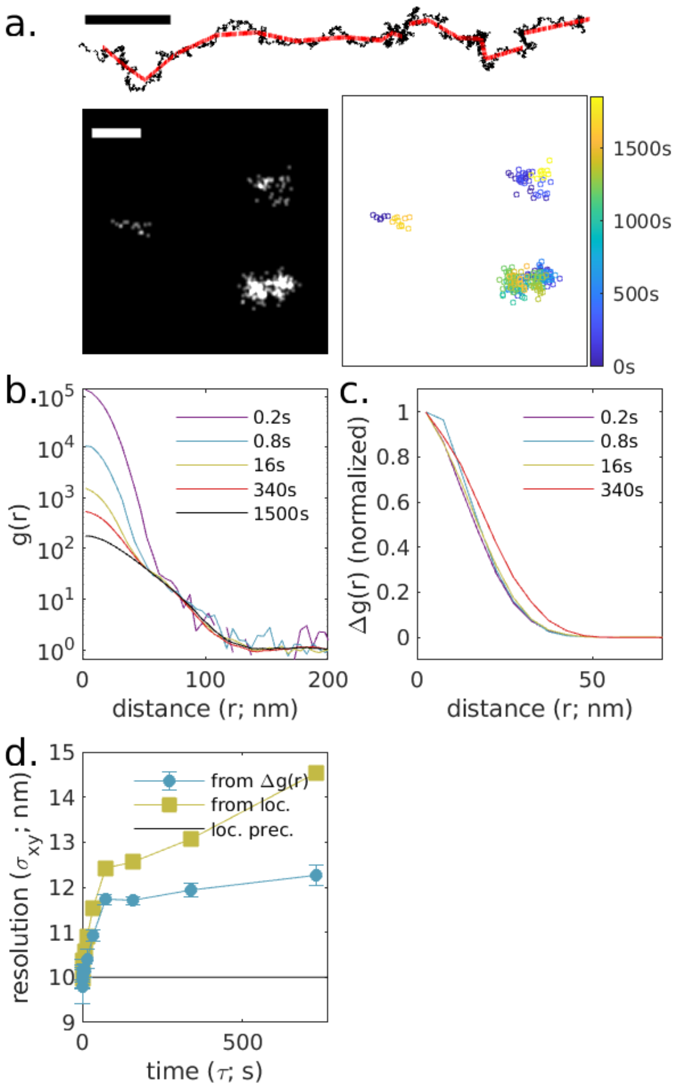


**Supplementary Figure S3: Simulation with drift and drift correction alongside single molecule motions.** (a.) The simulation from Fig 1 with applied drift (black) and drift correction (red) as shown in the trajectory above as well as single molecule diffusion with D=1nm$^2$/sec. Reconstructed image (left) and scatterplot of localizations with color representing the observation time (right) for a small subset of the simulated plane. Scale-bar is 100nm. (b.) Auto-correlations as a function of displacement, $g(r,\tau)$, tabulated from simulations for time-interval windows centered at the values shown. (c.) $\Delta g(r,\tau) = g(r,\tau) - g(r,\tau = 1500s)$ for the examples shown in b. (d.) $\Delta g(r,\tau)$ are fit to $\Delta g(r,\tau) \propto \exp\left\{-r^2 / 4\sigma_{xy}^2\right\}$ to extract out the resolution in each lateral dimension (from $\Delta g(r)$). The resolution estimated from $\Delta g(r,\tau)$ varies with time-interval, and is systematically narrower than the point spread function measured by grouping localizations with molecules ((from loc.). This is due to the distortion effect demonstrated in Fig S1 and is characterized by a $\sigma_{xy}$ that increases with $\tau$.

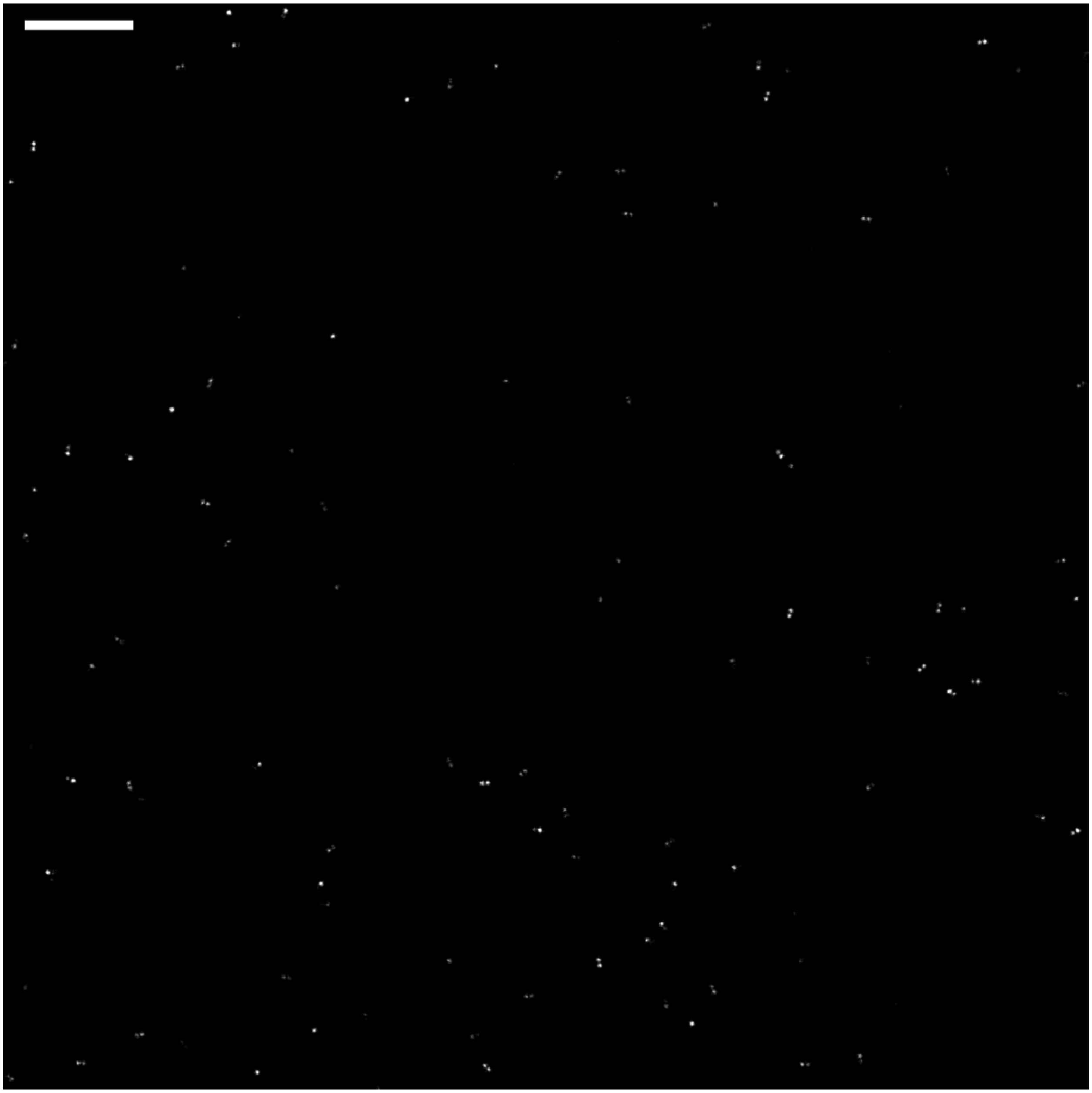

**Supplementary Figure S4.** 10µm by 10µm region showing simulated localizations from Figs 1-2. The full simulated area was 40µm by 40µm. Scale bar is 1µm.

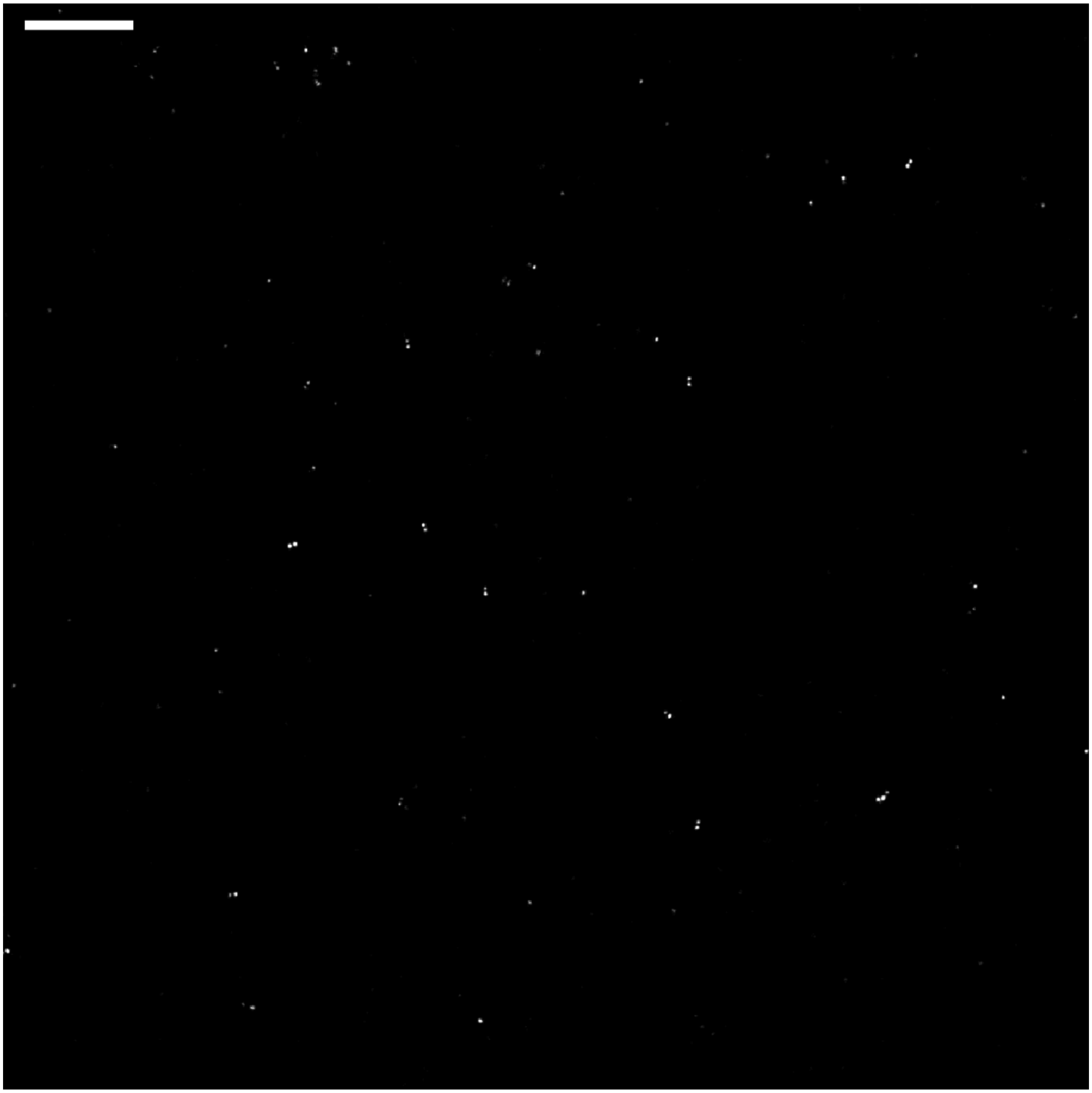

**Supplementary Figure S5.** 10µm by 10µm region showing DNA origami rulers analyzed in Fig 3. The full imaged area was 40µm by 40µm. Scale bar is 1µm.

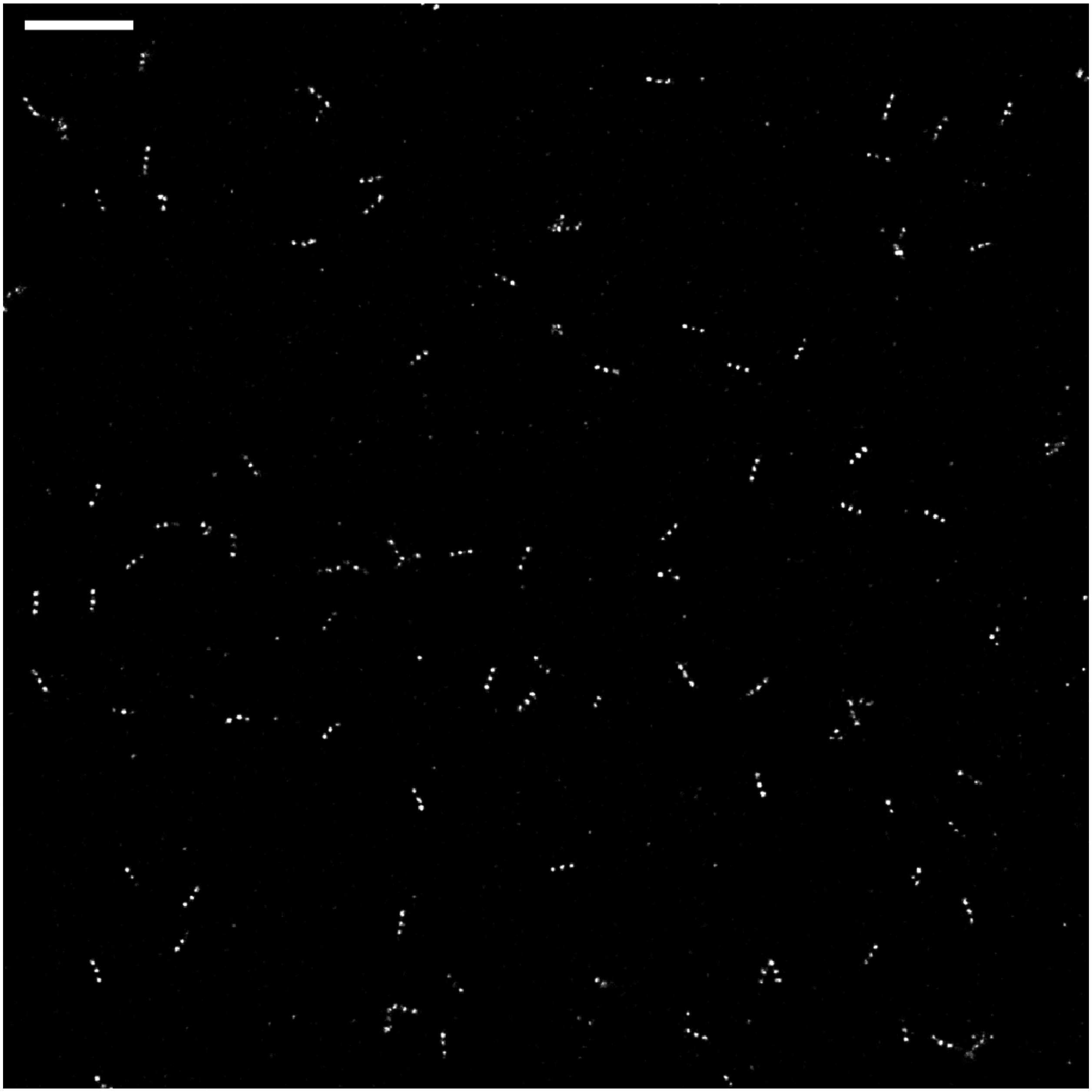

**Supplementary Figure S6.** 10µm by 10µm region showing DNA origami rulers analyzed in Fig 4. The full imaged area was 40µm by 40µm. Scale bar is 1µm.

**Supplementary Note: Derivation of spacetime pair correlation function estimator, and related computations.**

$N$ localizations $\mathbf{u}_i = (\vec{r}_i, t_i) = (x_i, y_i, t_i),\ i = 1, \ldots, N$ are observed on a spatial window (region of interest/ROI) $W$ during a temporal window $T$. This set of points is considered as a realization of a space-time point process $X$, so that we may define a (first-order) density $\rho(\mathbf{u}) = \rho(\vec{r}, t)$ notionally as

$$\frac{\text{Expected \# of points in area } d\vec{r} \text{ and time-interval } dt \text{ around } (\vec{r}, t)}{d\vec{r} \cdot dt}$$

Or more formally by the following:

$$\mathrm{E} \sum_{\mathbf{u} \in X \cap W \times T} 1[\mathbf{u} \in A] = \int_A \rho(\mathbf{u}) \mathrm{d}\mathbf{u} \tag{0.1}$$

For any set $A \subset W \times T$, where $1[\cdot]$ is an indicator function, taking the value 1 when its argument is true, and 0 otherwise. For the purposes of this paper, we assume that $\rho = \rho(t)$ is constant in space but may vary in time, e.g. due to bleaching of the fluorophores of the sample.

Further, define the second-order density $\rho^{(2)}(\mathbf{u}_1, \mathbf{u}_2)$ notionally by

$$\frac{\text{Expected \# of pairs of points in the } d\vec{r} \cdot dt \text{ neighborhoods of } \mathbf{u}_1 \text{ and } \mathbf{u}_2 \text{ respectively}}{(d\vec{r} \cdot dt)^2}$$

Or more formally

$$\mathrm{E} \sum_{\mathbf{u}_1, \mathbf{u}_2 \in X \cap W \times T}^{\neq} 1[\mathbf{u}_1 \in A \text{ and } \mathbf{u}_2 \in B] = \int_A \int_B \rho^{(2)}(\mathbf{u}_1, \mathbf{u}_2) \mathrm{d}\mathbf{u}_2 \mathrm{d}\mathbf{u}_1 \tag{0.2}$$

Now $\rho^{(2)}$ describes the second-order properties of $X$, for example attraction or repulsion between points. It is convenient to normalize $\rho^{(2)}$ so that it is dimensionless and easier to interpret. To that end, define the pair autocorrelation function $g(\mathbf{u}_1, \mathbf{u}_2)$:

$$g(\mathbf{u}_1, \mathbf{u}_2) = \frac{\rho^{(2)}(\mathbf{u}_1, \mathbf{u}_2)}{\rho(\mathbf{u}_1, \mathbf{u}_2)} \tag{0.3}$$

Loosely, the pair autocorrelation function is the ratio of the actual probability of finding points at both $\mathbf{u}_1$ and $\mathbf{u}_2$ to the hypothetical probability under the assumption that $\mathbf{u}_1$ and $\mathbf{u}_2$ are independent. We typically assume that $g$ is translation invariant in both space and time, and often further assume that it is rotationally invariant in space, so that it only depends on the separation of u1 and u2 in space and time, and we may write $g(\mathbf{u}_1, \mathbf{u}_2) = g(\|\vec{r}_2 - \vec{r}_1\|, t_2 - t_1)$.

We estimate $g$ using the standard kernel-based framework as laid out in e.g. (1, 2). Specifically, we use a box kernel with bandwidth $\delta_r$ in space and $\delta_t$ in time, and an isotropic edge-correction in space, and a density correction for the temporal edge correction, following the approach of (3). Briefly, consider the family of estimators for $g(r,\tau)$ given by:

$$\hat{g}(r,\tau) := \frac{1}{\gamma_{\text{sp.}}(r)\gamma_t(\tau)} \sum_{\mathbf{u}_i,\mathbf{u}_j \in X \cap W \times T}^{\neq} 1[\left|\left\|\vec{r}_j - \vec{r}_i\right\| - r\right| < \delta_r/2, \left|t_j - t_i - \tau\right| < \delta_t/2] \tag{0.4}$$

We wish to derive functions $\gamma_{\text{sp.}}$ and $\gamma_t$ such that the resulting estimator is unbiased. The expectation value of the sum in the above expression can be determined from an appropriate Campbell's theorem:

$$\begin{aligned}
\mathrm{E}\hat{g}(r,\tau) &= \frac{1}{\gamma_{\text{sp.}}\gamma_{\text{t}}} \int_{W\times T}\int_{W\times T} \rho^{(2)}(\mathbf{u}_1,\mathbf{u}_2) 1[\left|\left\|\vec{r}_2 - \vec{r}_1\right\| - r\right| < \delta_r/2, \left|t_2 - t_1 - \tau\right| < \delta_t/2] d\mathbf{u}_1 d\mathbf{u}_2 \\
&= \frac{1}{\gamma_{\text{sp.}}\gamma_{\text{t}}} \int_{W\times T}\int_{W\times T} g(\left\|\vec{r}_2 - \vec{r}_1\right\|, t_2 - t_1)\rho(t_1)\rho(t_2) 1[\left|\left\|\vec{r}_2 - \vec{r}_1\right\| - r\right| < \delta_r/2] 1[\left|t_2 - t_1 - \tau\right| < \delta_t/2] d\mathbf{u}_1 d\mathbf{u}_2 \\
&\approx \frac{g(r,\tau)}{\gamma_{\text{sp.}}\gamma_t} \int_W\int_W 1[\left|\left\|\vec{r}_2 - \vec{r}_1\right\| - r\right| < \delta_r/2] d\vec{r}_1 d\vec{r}_2 \int_T\int_T \rho(t_2)\rho(t_1) 1[\left|t_2 - t_1 - \tau\right| < \delta_t/2] dt_1 dt_2
\end{aligned}$$

Where the approximation in line 3 is due to the assumption that $g(r,\tau)$ is almost constant within $\delta_r/2$ in space and $\delta_t/2$ in time.

From the above derivation, it follows that $\hat{g}(r,\tau)$ is unbiased for the choices

$$\gamma_{\text{sp.}}(r) = \int_W\int_W 1[\left|\left\|\vec{r}_2 - \vec{r}_1\right\| - r\right| < \delta_r/2] d\vec{r}_2 d\vec{r}_1$$

$$\gamma_t(\tau) = \int_T\int_T \rho(t_1)\rho(t_2) 1[\left|t_2 - t_1 - \tau\right| < \delta_t/2] dt_2 dt_1$$

For computational considerations, we make further approximations on $\gamma_{\text{sp.}}$:

$$
\begin{aligned}
\gamma_{\text{sp.}}(r) &= \int_W \int 1[\left|\|\vec{h}\| - r\right| < \delta_r / 2] 1[\vec{r}_1 + \vec{h} \in W] d\vec{h} d\vec{r}_1 \\
&= \int_W \int_0^{2\pi} \int 1[|h - r| < \delta_r / 2] 1[\vec{r}_1 + (h\cos\theta, h\sin\theta) \in W] h dh d\theta d\vec{r}_1 \\
&= \int_W \int_{r-\delta_r/2}^{r+\delta_r/2} h \int_0^{2\pi} 1[\vec{r}_1 + (h\cos\theta, h\sin\theta) \in W] d\theta dh d\vec{r}_1 \\
&\approx r\delta_r \int_W \int_0^{2\pi} 1[\vec{r}_1 + (r\cos\theta, r\sin\theta) \in W] d\theta d\vec{r}_1 \\
&= r\delta_r \int_0^{2\pi} \left| W \cap W_{-(r\cos\theta, r\sin\theta)} \right| d\theta
\end{aligned}
$$

where $|A|$ indicates the area of the set $A$, and $A_{\vec{h}}$ indicates the translation of the set $A$ by the vector $\vec{h}$. The first line is a change of variables to $\vec{h} = \vec{r}_2 - \vec{r}_1$, with the extra indicator functions reflecting the integration bounds on $\vec{r}_2$, followed by a change to polar coordinates for $\vec{h}$. The approximation in the fourth line is justified by the fact that angular integral varies slowly with $h$, so the radial part of the integral can be approximately separated.

For the purposes of our matlab code, we represent the spatial window/ROI $W$ as a polygon with vertices `[mask.x(i),mask.y(i)]`. We translate the ROI by a vector `[hx,hy]` by simply adding `hx` and `hy` to `mask.x` and `mask.y`, respectively. Matlab provides functions `polybool` (to compute the intersection $W \cap W_{-\vec{h}}$, and `polyarea` to compute the area of the resulting polygon. It remains to complete the angular integral, which we compute by discretizing theta into 32 equally spaced points

$$
\gamma_{\text{sp.}}(r) = \frac{2\pi r\delta_r}{32} \sum_{i=1}^{32} \left| W \cap W_{-(r\cos\theta_i, r\sin\theta_i)} \right|, \qquad \theta_i = \frac{2\pi i}{32}
$$


1. Illian, J., A. Pentinen, H. Stoyan, and D. Stoyan. 2008. Statistical analysis and modelling of spatial point patterns. Chichester, England ; Hoboken, NJ: John Wiley.

2. Diggle, P. 2014. Statistical analysis of spatial and spatio-temporal point patterns. Boca Raton: CRC Press.

3. Shaw, T., J. Møller, and R. Waagepetersen. 2020. Globally intensity-reweighted estimators for $K$- and pair correlation functions. *ArXiv200400527 Stat*.
